# Localized estimation of electromagnetic sources underlying event-related fields using recurrent neural networks

**DOI:** 10.1101/2023.04.23.537963

**Authors:** Jamie A. O’Reilly, Judy D. Zhu, Paul F. Sowman

## Abstract

**Objective:** To use a recurrent neural network (RNN) to reconstruct neural activity responsible for generating noninvasively measured electromagnetic signals.

**Approach:** Output weights of an RNN were fixed as the lead field matrix from volumetric source space computed using the boundary element method with co-registered structural magnetic resonance images and magnetoencephalography (MEG). Initially, the network was trained to minimize mean-squared-error loss between its outputs and MEG signals, causing activations in the penultimate layer to converge towards putative neural source activations. Subsequently, L1 regularization was applied to the final hidden layer, and the model was fine-tuned, causing it to favour more focused activations. Estimated source signals were then obtained from the outputs of the last hidden layer. We developed and validated this approach with simulations before applying it to real MEG data, comparing performance with three existing methods: beamformers, minimum-norm estimate, and dynamical statistical parametric mapping.

**Main results:** The proposed method had higher output signal-to-noise ratios than the others and comparable correlation and error between estimated and simulated sources. Reconstructed MEG signals were also equal or superior to the other methods in terms of their similarity to ground-truth. When applied to MEG data recorded during an auditory roving oddball experiment, source signals estimated with the RNN were generally consistent with expectations from the literature and qualitatively smoother and more reasonable-looking than estimates from the other methods.

**Significance:** This work builds on recent developments of RNNs for modelling event-related neural responses by incorporating biophysical constraints from the forward model, thus taking a significant step towards greater biological realism and introducing the possibility of exploring how input manipulations may influence localized neural activity.

## 1. Introduction

The “Holy Grail” of neuroimaging, is a non-invasive modality that can instantaneously and comprehensively observe neural activity at any location of the nervous system, ideally capable of distinguishing between signals reflecting different biochemical interactions. While the search for this technology is ongoing, contemporary neuroscientists face a trade-off between high spatial resolution, structural imaging modalities, such as X-ray computed tomography (CT) or magnetic resonance imaging (MRI), and high temporal resolution, functional neuroimaging modalities, such as electroencephalography (EEG) or magnetoencephalography (MEG). Functional MRI measures blood oxygenation signals with lower temporal resolution than M/EEG and do not directly reflect the brain’s bioelectric communication. To infer localized neural activity that unfolds over tens to hundreds of milliseconds, the scale required for detailed examination of sensory and cognitive processes, it is necessary to use either EEG or MEG in conjunction with structural imaging data. The magnetic fields measured with MEG suffer less attenuation and scattering than the electric fields measured with EEG as signals pass through the skull and scalp; hence MEG is considered to be more reliable for estimating source signals [1].

Localized source estimation generally involves source space, sensor space, a forward model, and an inverse model [2]. The source space defines the locations of prospective neural sources within the brain. The sensor space defines the locations and geometry of the sensors used to acquire signals from outside the head. Source and sensor spaces are co-registered to the same coordinate scheme. The forward model is built from physical assumptions about tissue morphology, conduction velocity, signal acquisition modality, and co-registered source and sensor space geometry. This information is used to calculate weightings that estimate the contribution of each source location (in x-, y- and z-directions) to each sensor. These provide a mapping from source signals to sensor signals, called the lead field matrix. The inverse model is an operation that performs the inverse transformation from sensor signals to source signals. However, the lead field matrix is non-invertible, and for real MEG signals we cannot ascertain the true underlying source signals, making this an ill-posed problem.

Several methods have been developed to tackle this inverse modelling problem, including dipole fitting, adaptive spatial filtering, and minimum-norm optimisation approaches [2]. Synchronous ionic currents through apical dendrites of cortical pyramidal cells are thought to produce the majority of electromagnetic signals that can be detected from outside the head using current technology [3]. These can be modelled as current dipoles with a magnitude and orientation. In dipole fitting, estimating the location, orientation and magnitude of one or several current dipoles is treated as an optimization problem, with the objective of explaining the observed pattern of sensor signals. This typically involves iterative optimization that arrives at a non-unique solution that can vary widely depending on initial conditions. Adaptive spatial filtering (e.g., beamformers) derives a set of optimized spatial filters that minimize covariance among source estimates; these are applied to distributed source spaces that can include thousands of fixed source locations [4,5]. This approach underestimates highly covariant sources, and does not consider the degree to which estimated source signals explain observed sensor signals. Minimum norm estimates (MNE) are also applied to distributed source spaces, optimizing amplitudes and orientations to minimize the difference between observed sensor signals and those reconstructed from estimated source signals, subject to additional constraints such as minimizing the L2 norm of estimated source amplitudes [2,6]. An extension to the MNE approach, called dynamic statistical parametric mapping (dSPM), involves scaling the amplitudes of source estimates based on the noise covariance matrix at each source location [7]. Whilst beamforming, MNE and dSPM methods are widely used, all have limitations that can be challenging to quantify due to the lack of ground-truth for neural signals underlying genuine MEG data. Simulations can be used to evaluate source estimation methods [8], although it may be prudent to apply multiple methods to real data given the uncertainty.

Recurrent neural networks (RNNs) have recently been developed for modelling auditory-evoked epidural field potentials from mice [9,10], auditory event-related potentials from humans [11], and visual event-related potentials from humans [12,13]. These models learn transformations between input representations of physical events and average event-related neural signals, which can be probed with simulations and examined using data analysis techniques. By fitting labels from evoked neural activity in a supervised learning paradigm, hidden units of the RNN produce signals that can be interpreted as activity of putative (non-localized) sources [9-13]. In the present study, we expand this toolset by incorporating the lead field matrix into the network architecture to impose spatial priors on estimated source signals from the RNN. We tested this method on simulations then applied it to real MEG signals and compared the results with some existing source estimation techniques. We acquired the MEG data from a roving auditory oddball paradigm experiment designed to elicit the magnetic counterpart of mismatch negativity (MMNm), which is considered to arise from neural sources located bilaterally in the temporal lobes and in the frontal lobe [14].

## 2. Materials and methods

### 2.1. Magnetoencephalography

To collect data for this study we conducted a roving auditory oddball paradigm experiment with a single subject (author J.D.Z.) at the KIT-Macquarie Brain Research Laboratory in Sydney, Australia, using previously described equipment [15]. The experimental protocol was reviewed and approved by the Human Research Ethics Committee at Macquarie University. Recordings were made inside a magnetically shielded room (Fujihara Co. Ltd., Tokyo, Japan) using an MEG system with 160 axial gradiometers covering the whole head (Model PQ1160R-N2; Kanazawa Institute of Technology, Kanazawa, Japan). Auditory stimuli were sequenced using custom Matlab (version R2017b; The MathWorks Inc., Natick, MA, USA) scripts and delivered binaurally via tubal insert earphones (Etymotic ER30; Etymotic Research, Inc., Elk Grove Village, IL, USA).

Before recording MEG, subject head shape information was obtained using a digitizer (Fastrak; Polhemus, Colchester, VA, USA). Nasion (N) and bilateral preauricular area (LPA and RPA) locations were also recorded using the digitizer. The subject wore a tight-fitting elastic cap with five head position indicator coils attached, which were recorded by the digitizer and then used to monitor head position within the MEG sensor array during the recording session. Head position was found to deviate <5 mm.

Continuous MEG signals were recorded at 1000 Hz sampling rate with online 0.03 to 200 Hz band-pass filtering. An empty room recording was also conducted and this was used to reduce environmental noise by applying the time-shifted principal component analysis algorithm [16] implemented in MEG Laboratory software (Yokogawa Electric and Eagle Technology) with 10 s block width and three shifts. Signals were resampled offline to 200 Hz.

### 2.2. Roving oddball sequence and data extraction

A version of the roving oddball paradigm [17,18] was used in this experiment, which consisted of 224 trains of between 6 and 10 repetitions of one of seven different frequency tones (500, 550, 600, 650, 700, 750, 800 Hz). These tones were 70 ms sinusoids with 5 ms rise/fall ramps, calibrated to reach 80 dB sound pressure level. The inter-stimulus interval was 500 ms. Participants focused on a fixation cross throughout the task. Responses to the first and last stimuli in this sequence were discarded. From the remaining stimulus trains, standard trials were extracted from 0.1 s before to 0.4 s after the final tone in each tone-train, while deviant trials were extracted from 0.1 s before to 0.4 s after the first tone in each train. This resulted in 223 standard and 223 deviant trials, each with 100 time-samples, recorded from 160 MEG sensors, producing a data matrix sized (446, 100, 160).

Signal-to-noise ratio (SNR) of MEG single trials was calculated using equation (1), where Signal_RMS_ was the root-mean-squared (RMS) value computed from the post-stimulus time-window (>0 s) and Noise_RMS_ was calculated from the pre-stimulus time window (<0 s); calculated to be 2.79 dB. Event-related field (ERF) waveforms were produced by averaging standard and deviant trials, producing matrices sized (100, 160) for standard and deviant ERFs. These had SNRs of 4.89 dB and 9.74 dB, respectively. We computed the RMS of their corresponding ERF waveforms across all channels to visualize the overall brain response elicited by standard and deviant conditions.

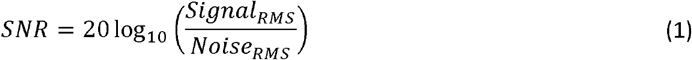

### 2.3. Forward model

Code for computing the forward model is available from the KIT-Macquarie Brain Research (MEG) Laboratory GitHub repository (https://github.com/Macquarie-MEG-Research/). A structural (T1-weighted) MRI scan of the subject was manually co-registered with head shape information from the MEG data using a graphical user interface [19]. Head position indicators on an elastic cap worn during the recording session were used to align the MEG sensors into the same space. Thus, the MRI voxels, MEG sensor positions, and source locations (Figure 1) were transformed into a common frame of reference.

**Figure 1.**
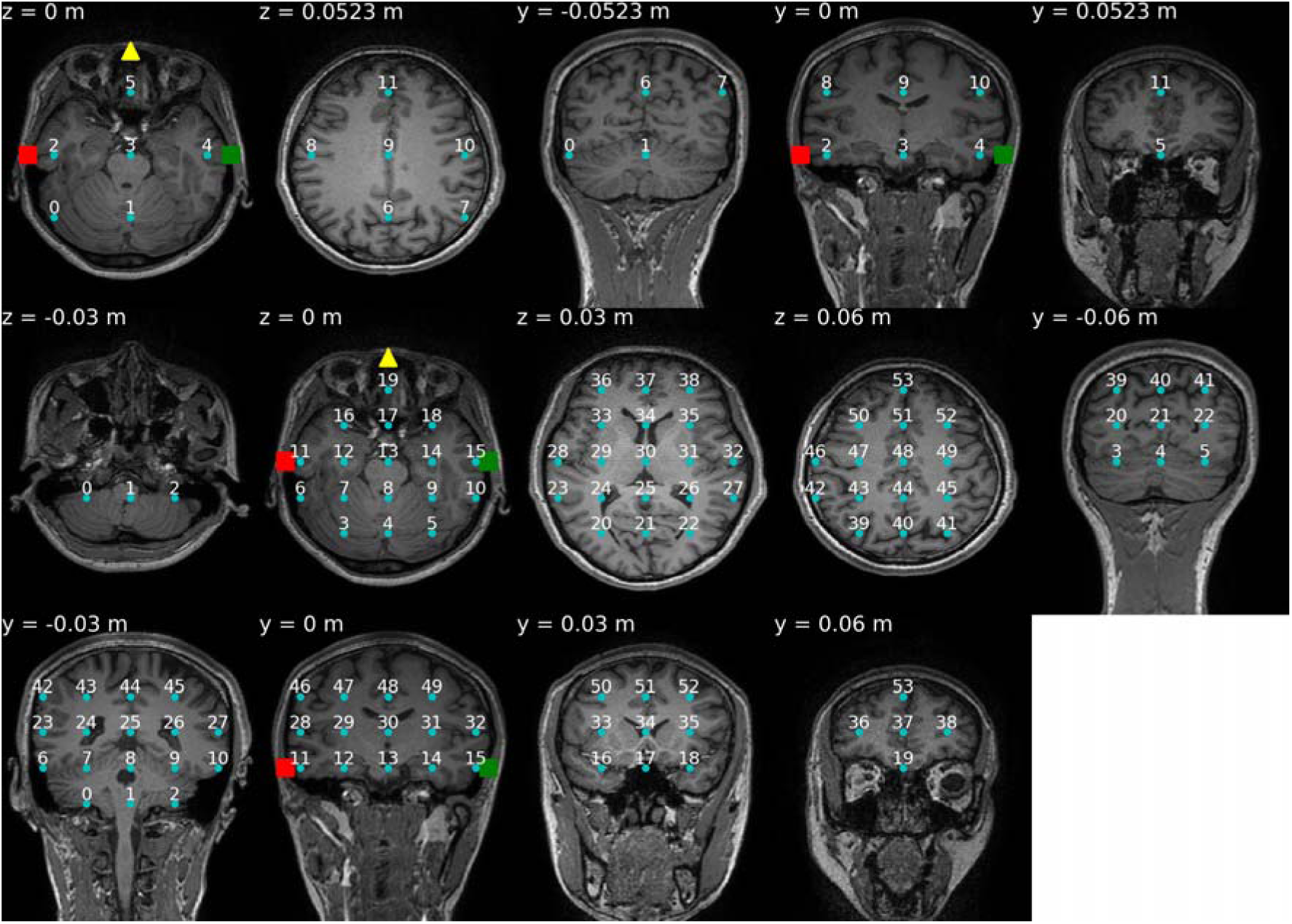
Volumetric source spaces co-registered with structural neuroimaging data. The top row shows 12-source source space produced with 0.0523 m grid spacing. The middle and bottom rows show 54-source source space produced with 0.03 m grid spacing. Slice location is indicated in the upper-left on each panel; z coordinates are given for slices in the xy plane, and y coordinates are given for slices in the xz plane. Source indices are noted above each source location (blue circle). The nasion is indicated with a yellow triangle, and left and right preauricular areas are indicated with red and green squares.

A boundary element method (BEM) volume conduction model was constructed from the inner skull segment and used as the forward model to define two volumetric source spaces, each with a different number of sources (*n*_*src*_). A 12-source space (*n*_*src*_ = 12) was computed with 5.23 cm grid spacing, and a 54-source space (*n*_*src*_ = 54) was computed with 3 cm grid spacing. While the former is sparser, even the 54-source source space may be considered sparse relative to some distributed source models [20]. Nevertheless, these were selected as a proof-of-concept for evaluating the proposed method of localized source estimation. Lead field matrices (*L*) of resulting forward models were sized (160, 3*n*_*src*_); with contributions to MEG sensor measurements of each source in x-, y-, and z-directions weighted separately. Transformation of source signals into corresponding MEG sensor signals via the lead field matrix is achieved as in equation (2), where the sources matrix is sized (100, 3*n*_*src*_) and the MEG signal matrix is sized (100, 160). However, the inverse operation is ill-defined as there is no unique solution to transform MEG signals back into corresponding source signals. Also, lacking ground-truth measurements of the source signals underlying real MEG data, evaluating source reconstruction methods typically requires synthetic MEG obtained from simulated source signals using the lead field matrix.

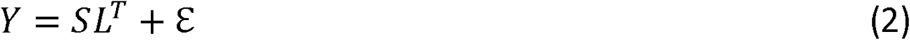

Where *Y* ∈ ℝ^T×M^ is an MEG signal matrix with T = 100 time-samples and M= 160 sensors, *S*∈ ℝ^T×N^ is a source signal matrix with T time-samples and *N* = 3*n*_*src*_ source components, *L* ∈ ℝ^M×N^ is the lead field matrix, and ε ∈ ℝ^T×M^ is a matrix of extraneous noise signals generated by biological and non-biological sources. Because the inverse problem is ill-posed, the exact value of *S* is theoretically unknowable.

### 2.4. Simulations

Simulated data was designed to mimic the structure of real MEG data extracted from recordings made during the roving auditory oddball sequence. We selected two source locations to simulate “auditory” activations. These were located bilaterally in the temporal lobes at source indices 2 and 4 for 12-source source space, and 11 and 15 for 54-source source space. Another source location was selected for the theoretical “cognitive” component of the MMNm response, located at source index 11 for 12-source source space and 53 for 54-source source space. These sources are identified in Figure 1.

Standard and deviant conditions were simulated for 226 trials each, matching the structure of the real data. Each trial was considered to span the same time range as those extracted from MEG recordings. To simulate auditory responses at preselected temporal source locations, two cycles of a 10 Hz sinusoid were generated, starting at 0 s and ending at 0.2 s; for standard trials the peak amplitude was set to 10 nA, and for deviant trials this was 15 nA. Different amplitudes of these simulated auditory responses were selected to imitate sensory adaptation differences between standard and deviant responses which are thought to influence auditory responses elicited during the roving oddball sequence [18]. A Hann-windowed signal (onset 0.1 s and offset 0.3 s; peak amplitude of 10 nA) was generated to simulate the cognitive component of the MMNm response. This signal was only applied to the preselected frontal source location in deviant trials, representing the cognitive processes considered to underlie MMNm [21]. Pink (1/f) noise was generated as described previously [22] and superimposed upon these simulated source activations. Simulated MEG signals were derived from these simulated source activations by multiplying them with the lead-field matrix of the forward model according to equation (3). Source estimates were derived from simulated MEG signals using RNN and beamformer methods, and subsequently compared with the ground-truth simulated source activations using Pearson’s correlation coefficient (r) and mean-squared error (MSE).

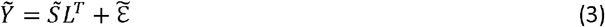

Where 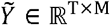 is a simulated MEG signal matrix, 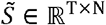 is a simulated source signal matrix, and 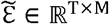 is a simulated noise signal matrix. Generating synthetic data in this manner allows us to evaluate methods of source signal estimation where the ground-truth 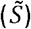 is known. Here the tilde accent is used to distinguish simulated source and MEG signals from their real counterparts.

Two sets of simulations were produced: one with all source activations equal in magnitude across three spatial dimensions, producing non-optimally oriented resultant vectors, and another with each source activation specified to reflect its optimal orientation in three dimensions. The optimal orientation for each source was determined by squaring the lead-field matrix then summing x-, y-, and z-components (160 of each, reflecting the number of MEG channels) and dividing the resulting vector by its magnitude to produce a three-element unit vector. This unit vector points in the direction of maximum sensitivity of MEG to activity generated at the source location, based on the geometry and theoretical assumptions asserted by the forward model. Non-optimal and optimal orientations for each source location in 12- and 54-source source spaces are depicted in Supplementary Figure S1. All trials for non-optimally and optimally oriented simulated source activations for 12- and 54-source source spaces are plotted in Supplementary Figures S2-S5.

Non-optimally oriented and optimally oriented simulated source activations had SNR of 5.5 dB for 12-source source space and 1.96 dB for 54-source source space; calculated over all trials using equation (1). Corresponding simulated MEG for non-optimally oriented sources had SNRs of 3.95 dB and 1.92 dB for 12- and 54-source source spaces; optimally oriented sources had SNRs of 5.91 dB and 3.08 dB for 12 and 54 sources. Difference between ERFs produced by non-optimally and optimally oriented 12- and 54-source simulations reflects how the lead-field matrix weights sources according to their direction.

### 2.5. RNN source estimation

The model architecture and supervised training scheme are illustrated in Figure 2. Input arrays were shaped 100 (time-samples) by 2 (event channels). Each event channel of the input arrays was a unit step pulse function for the respective event (i.e., either the standard or deviant auditory event). The output labels used for supervised learning consisted of single-trial MEG data recorded during respective events, each shaped 100 (time-samples) by 160 (MEG sensors), scaled up by a factor of 10^13^. As described above, 223 trials of standard and deviant events from the roving oddball sequence were extracted. Associating each input representation with multiple output labels and optimizing for the mean-squared-error loss function produces model outputs that converge towards the average of those labels, allowing the RNN to reproduce ERFs computed from those single-trial labels.

**Figure 2.**
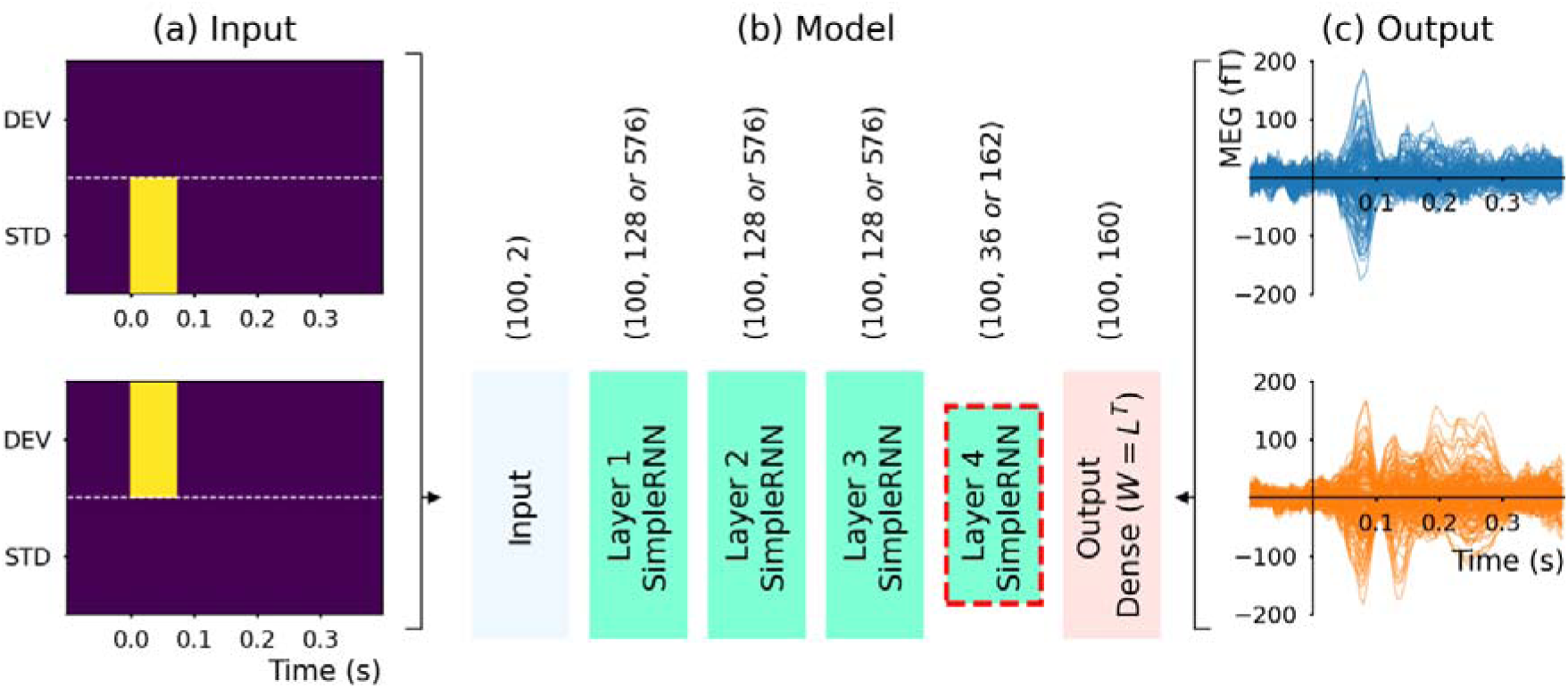
Localized source estimation using a recurrent neural network. Input representations (a) of standard and deviant trials were mapped to output labels (c) from 160 MEG channels. This produces a one-to-many mapping, associating each input with multiple output labels (from 223 trials), causing model outputs to converge towards the average of those labels by minimizing mean-squared error loss. The model (b) consisted of an input layer followed by four hidden recurrent layers (SimpleRNN TensorFlow layer class) and a fully-connected output layer (Dense layer class). The data shape out from each layer is indicated above it, with 100 time-samples throughout and the number of artificial neurons in layers 1-4 varying according to the source space (specified as 12-source *or* 54-source values). The output layer had 160 units to equal the number of MEG channels used as labels, and its weights were fixed as the lead field matrix from the forward model. The number units in layer 4 was determined by the number of sources multiplied by three spatial directions (i.e., 36 for 12 sources *or* 162 for 54 sources). Layer 4 is the source estimation layer because its outputs feed into the final layer that effectively performs equation (4) to estimate MEG signals. The model was trained in two phases: (1) without any constraints beyond minimizing mean-squared error loss, and (2) with L1 activity regularization applied and bias removed from layer 4 units, causing the model to minimize loss with greater efficiency in terms of source activations.

The model architecture is illustrated in Figure 2b. It comprised an input, four recurrent (SimpleRNN) hidden layers, the first three with rectified linear unit (ReLU) activation function and the fourth with linear activation, and a fully-connected (Dense) output layer with linear activation. The input layer was defined to accept input arrays shaped (100, 2), suitable for the aforementioned input representations. The output layer was defined to generate outputs shaped (100, 160), which were compared directly with output labels during training. The output layer did not use a bias, and its weights were fixed as the lead field matrix multiplied by a factor of 10^6^, such that the output layer effectively implemented equation (4).

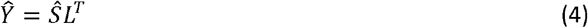

Where 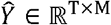 an estimated MEG signal matrix and 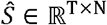 is an estimated source signal matrix. Here the wide-hat accent denotes estimated MEG and source signals, to distinguish them from real and simulated counterparts. The source estimation method must mitigate the influence of extraneous noise.

Numbers of units in hidden layers were selected based on the forward model’s source space and resulting lead field matrix. The number of units in the final hidden layer (layer 4; the source estimation layer) was either 36 (12 × 3 directions) for the 12-source space, or 162 (54 × 3 directions) for the 54-source space. Source signal components in x, y and z directions were thus treated independently. The three preceding hidden layers were designed to have either 128 units for 12 sources, or 576 units for 54 sources. In preliminary tests, more units in hidden layers preceding the source estimation layer were found to improve source estimation performance up to a point, but increasing them above a ratio of 3.56 yielded no further improvements.

Output from the input layer can be denoted as in equation (5).

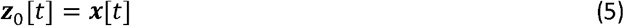

Where ***x***[*t*] ∈ {0, 1}^1 × 2^ is a vector containing the input signal for time-sample *t* ∈ {1: 100}; thus *X* ∈ {0, 1}^100 × 2^. At each time-sample, the possible states of ***x***[*t*] were either [0, 0], [1, 0] (standard), or [0, 1] (deviant), as illustrated in Figure 2a.

The formula implemented by subsequent recurrent layers can then be formulated as in equation (6).

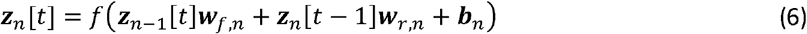

Where ***z***_*n*_[*t*] ∈ ℝ^1 × *P*^ (*P* ∈ {128, 36, 576, 162}) is the output from units in layer *n* ∈ {1, 2, 3, 4} at time-sample *t, f* (·) is the activation function (*ReLU* for layers 1-3, *linear* for layer 4), ***z***_*n* − 1_ [*t*] ∈ ℝ^1 × *Q*^ (*Q* ∈ {2, 128, 576}) is the output from units in layer *n* − 1 at time sample *t*, ***z***_*n*_[*t* − 1] ∈ ℝ^1 × *P*^ is the output from layer *n* units at time-sample *t* − 1 (initialised as zeros for *t* = 1), ***w***_*f, n*_ ∈ ℝ^*Q* × *P*^ are the forward weights for layer *n*, ***w***_*r, n*_ ∈ ℝ^*P* × *P*^ are the recurrent weights for layer *n*, and ***b***_*n*_ ∈ ℝ^*P*^ is the bias vector for units in layer *n*.

For transfer learning, the bias vector was removed from recurrent layer 4 (i.e., the source estimation layer), changing equation (6) into equation (7).

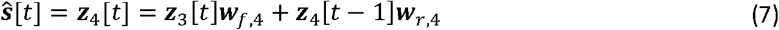

Where 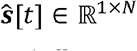 (*N* ∈ {36, 162}) is a vector of estimated source signals at time-sample *t*, ***z***_3_ [*t*] ∈ ℝ^1 × *K*^ (*K* ∈ {128, 576}) are the outputs from layer 3, ***w***_*f*, 4_ ∈ ℝ^*K* × *N*^ are the forward weights, ***z***_4_ [*t* − 1] ∈ ℝ^1 × *N*^ are the outputs from layer 4 and time-sample *t* − 1, and ***w***_*r*, 4_ ∈ ℝ^*N* × *N*^ are the recurrent weights for layer 4. No activation function was applied, thus facilitating positive and negative estimated source amplitudes.

The output layer performed equation (8).

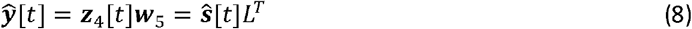

Where 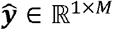 (*M* = 160 sensors) is the predicted output at time-sample *t*, and ***z***_4_ [*t*] ∈ ℝ^1 × *N*^ are the outputs from layer 4 or the estimated source signals, and ***w***_5_ = *L*^*T*^ ∈ ℝ^N × *M*^ are the forward weights of the output layer (layer 5) which were fixed as the transposed lead field matrix. When performed over all of the time-samples, the output layer effectively enacts equation (4).

Models were trained in two phases. In step 1, they were trained to minimize mean-squared-error loss between model outputs and single-trial MEG labels, calculated as in equation (9).

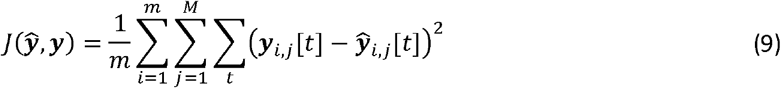

Where *J* (·) denotes the loss function, *m* is the number of training instances in each batch, ***y*** _*i, j*_ [*t*] are the labels and 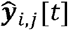 are the model predictions from MEG sensor index *j*. The loss is calculated over all time-samples and fed-back to update the model weights and biases using the backpropagation through time training (BPTT) algorithm implemented in TensorFlow.

In training step 2, the bias vector was removed from layer 4 as noted in equation (7), and L1 activity regularization was applied to layer 4 outputs, thus modifying the loss function in equation (10).

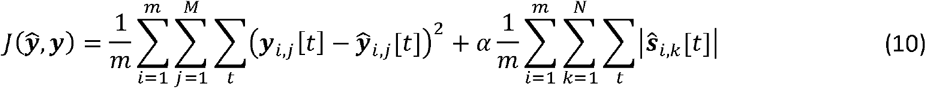

Where *α* = 0.0001 was a coefficient to control the penalty associated with the L1-norm of layer 4 activations, and 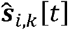 was the amplitude of estimated source signal *k* for training instance *i*.

Transfer learning from step 1 with the additional constraint of minimizing the L1 norm of layer 4 activations was used to derive source estimates. In preliminary tests, removing the layer 4 bias was necessary for producing reasonable source estimates after introducing L1 norm regularization. Substantial baseline deviations occurred if the bias was not removed and the model could not fit the output labels.

Optimization was performed in both training phases using BPTT with the adaptive moment estimation (Adam) optimizer. Default hyperparameters were used: learning rate 0.001, beta-1 0.9 and beta-2 0.999. Training was conducted for a maximum of 1000 epochs with batch size of 100; although this was stopped early if the loss failed to decrease for 50 consecutive epochs. After training, source estimates were obtained from the activations of layer 4 in response to input representations, multiplied by a factor of 10^−7^. Appropriately scaling output labels (10^13^), lead field matrix (10^6^) and layer 4 activations (10^−7^), mitigated potential training issues related to vanishing or exploding gradients [23], and estimated source amplitudes in the anticipated nano-Ampere (nA) range.

Model hidden layers had forward weights initialised using the Glorot uniform distribution [24], recurrent weights initialised from an orthogonal distribution [25], and biases initialised as zeros. The random seed for model initialisations was 0 for simulations. For real MEG data, five models were trained with random seeds 0 to 4 and the average of resulting source estimates was taken. Correlations between source estimates derived from different model initialisations were analysed as shown in Figure 5.

### 2.6. Other methods of source estimation

We compared source estimates generated from the RNN approach with the results from three other source estimation methods implemented in MNE-Python [19].

Linearly Constrained Minimum Variance (LCMV) spatial filtering weights, referred to as the beamformer (BF) method, were used for source estimation [4,26]. Unit-noise gain minimum BFs were computed with freely oriented sources, yielding vector current components in three separate directions. Noise covariance was computed from the pre-stimulus baseline window (−0.1 to 0 s), and data covariance was computed from the post-stimulus window (0 to 0.4 s) and whitened with 0.05 regularization. Spatial filters were computed from the average of all events (i.e., combined standard and deviant trials), then applied to separate ERFs to estimate patterns of source activity that could be responsible for generating standard and deviant ERFs. The BF approach is a parametric scanning method of source estimation that is known to exhibit relatively poor performance when source activations are highly correlated [2,27].

We also applied minimum norm estimate (MNE; Hämäläinen and Ilmoniemi, 1994) and dynamic statistical parametric mapping (dSPM; Dale et al., 2000) approaches. Both of these methods minimize squared difference between reconstructed and real ERFs and the L2-norm of estimated source activations. The dSPM method additionally includes dividing the source estimate amplitudes by the estimated noise at the respective source location [2,7]. Both MNE and dSPM methods yielded vector magnitudes for freely oriented source estimates by combining three direction components, using a depth prior of 0.8. When applying the inverse operator, the regularization parameter (lambda2) was set to 1/9. These methods produced identical ERFs, hence when comparing reconstructed ERFs in Figures 4 and 9 these are pooled together. Importantly, these methods are not known to suffer from the same limitations as the BF approach concerning highly covariant source activity.

### 2.7. Data Analysis

The vector magnitudes of the source estimates were compared with those from simulated source signals using Pearson’s correlation coefficient (r) and mean-squared error (MSE). Signal-to-noise ratios (SNRs) of source signals estimated by different methods were also computed. Reconstructed ERFs from sources estimated from simulated MEG data were also compared using correlation coefficient, MSE and SNR. These values are reported in Table 1.

**Table 1.** Comparison of source estimation methods applied to simulated source activations.

| Simulation | Estimation method | STD SRC (r, MSE) | DEV SRC (r, MSE) | SRC SNR (dB) | STD ERF (r, MSE) | DEV ERF (r, MSE) | ERF SNR (dB) |
| --- | --- | --- | --- | --- | --- | --- | --- |
| 12 sources, non-optimal orientation | BF | .284, 3.04 | 0.4, 6.96 | 19.9 | .83, 37.8 | .399, 71.7 | 12.9 |
|  | dSPM | .999, .198 | 1, .503 | 30.2 | 1, 1.13 | 1, 2.69 | 27.7 |
|  | MNE | .999, .079 | 1, .186 | 30.3 | 1, 1.13 | 1, 2.69 | 27.7 |
| | RNN | .922, .593 | .918, 1.54 | ∞ | 0.998, .462 | 1, .584 | ∞ |
| 12 sources, optimal orientation | BF | .23, 3.08 | .39, 7.03 | 19.6 | .927, 76.8 | .401, 144.0 | 17.7 |
|  | dSPM | .999, .198 | 1, .498 | 30.2 | 1, 2.11 | 1, 4.96 | 30.7 |
|  | MNE | .999, .0781 | 1, .186 | 30.2 | 1, 2.11 | 1, 4.96 | 30.7 |
| | RNN | .986, .15 | .994, .202 | ∞ | .998, .416 | .999, .702 | ∞ |
| 54 source, non-optimal orientation | BF | .284, .677 | .425, 1.53 | 13.9 | .743, 48.1 | .443, 94.9 | 13.2 |
|  | dSPM | .97, .135 | .974, .326 | 25 | .999, 5.26 | 1, 12.9 | 23.8 |
|  | MNE | .943, .0983 | .951, .237 | 24.5 | .999, 5.26 | 1, 12.9 | 23.8 |
| | RNN | .892, .261 | .898, .642 | ∞ | .998, 1.06 | .998, 1.62 | ∞ |
| 54 sources, optimal orientation | BF | .273, 0.68 | .421, 1.54 | 13.7 | .827, 92.7 | .465, 179.0 | 15.6 |
|  | dSPM | .992, .0805 | .994, .202 | 24.9 | 1, 8.64 | 1, 22.1 | 26.6 |
|  | MNE | .987, .0556 | .989, .14 | 24.2 | 1, 8.64 | 1, 22.1 | 26.6 |
| | RNN | .995, .0259 | .995, .0541 | ∞ | .999, 0.811 | .999, 0.832 | ∞ |

Abbreviations: simulated standard (STD), simulated deviant (DEV), source (SRC), event-related field (ERF), mean-squared error (MSE), signal-to-noise ratio (SNR). Correlation coefficients were calculated from the vector magnitudes of estimated and simulated sources in three directions, and from root-mean squared power across 160 channels for ERFs; all had $p < 0.001$ ; MSEs from ERFs were calculated after normalizing to fT range.

When applying source estimation methods to real data there is no ground-truth to compare with the results. As such, only SNR was calculated from the vector magnitudes of source signals estimated to underlie real MEG data. In addition, reconstructed ERFs produced by sources estimated from real MEG data were compared in terms of correlation, MSE and SNR. These measurements are reported in Table 2.

**Table 2.** Comparison of source estimation methods applied to real MEG data.

| Number of sources | Estimation method | SRC SNR (dB) | STD ERF (r, MSE) | DEV ERF (r, MSE) | ERF SNR (dB) |
| --- | --- | --- | --- | --- | --- |
| 12 | BF | 6.22 | .704, 201.0 | .811, 410.0 | 6.02 |
|  | dSPM | 8.43 | .975, 31.9 | .954, 53.9 | 9.21 |
|  | MNE | 8.26 | .975, 31.9 | .954, 53.9 | 9.21 |
|  | RNN | 14.3 | .984, 29.3 | .991, 18.0 | 12.4 |
| 54 | BF | 5.88 | .578, 60.6 | .778, 69.8 | 4.92 |
|  | dSPM | 5.96 | .99, 14.2 | .991, 11.7 | 8.87 |
|  | MNE | 5.65 | .99, 14.2 | .991, 11.7 | 8.87 |
|  | RNN | 14 | .981, 35.6 | .99, 30.9 | 12.7 |

Source estimates obtained from each method were ranked by SNR from highest to lowest, as an approximation of their relative importance. Source estimates were also ranked by correlation among methods from highest to lowest. For the RNN method only, each source was also ranked by setting the other sources to zero, then measuring correlation and MSE between real and reconstructed ERFs; this was repeated for each source individually, and they were then ranked (i) by correlation from highest to lowest, and (ii) by MSE from lowest to highest. The top ten sources from each of these ranking methods are reported in Table 3.

**Table 3.** Top ten ranked sources from different source estimation methods

| Source space | Estimation method | Ranking method | Top ten sources (1 to 10) |
| --- | --- | --- | --- |
| 12 | BF | SNR | 10, 4, 8, 3, 7, 11, 9, 1, 2, 5 |
|  | dSPM | SNR | 10, 8, 4, 9, 3, 6, 7, 11, 5, 2 |
|  | MNE | SNR | 10, 8, 4, 9, 3, 6, 7, 11, 5, 2 |
|  | RNN | SNR | 10, 3, 2, 4, 11, 1, 6, 5, 0, 9 |
|  | All | Correlation among estimation methods | 3, 8, 9, 10, 4, 7, 5, 2, 6, 11 |
|  | RNN | Correlation with reconstructed ERFs | 8, 6, 9, 11, 10, 1, 3, 5, 4, 7 |
|  | RNN | MSE with reconstructed ERFs | 8, 10, 4, 6, 7, 5, 9, 2, 3, 0 |
| 54 | BF | SNR | 22, 31, 35, 32, 15, 30, 17, 43, 28, 14 |
|  | dSPM | SNR | 32, 15, 27, 46, 17, 31, 14, 38, 52, 35 |
|  | MNE | SNR | 32, 15, 27, 46, 17, 31, 14, 38, 52, 35 |
|  | RNN | SNR | 52, 40, 15, 35, 49, 19, 32, 42, 46, 44 |
|  | All | Correlation among estimation methods | 32, 46, 31, 44, 14, 6, 35, 33, 30, 42 |
|  | RNN | Correlation with reconstructed ERFs | 32, 23, 28, 10, 27, 6, 11, 46, 36, 39 |
|  | RNN | MSE with reconstructed ERFs | 32, 46, 27, 23, 28, 15, 52, 41, 39, 42 |

SNR and correlation ranks are ordered from highest to lowest; error ranks are ordered from lowest to highest.

Another analysis of the RNN approach involved consecutively turning on sources from highest to lowest SNR and measuring correlation between real and reconstructed ERFs. These data are plotted in Figure 8 and illustrate the relative contributions that adding each source has on reproducing ERF waveforms.

### 2.8. Software

This work was conducted using Python 3 with MatPlotLib 3.5.2, MNE-Python 1.0.3, NumPy 1.22.4, SciPy 1.8.1, and TensorFlow 2.11.0. Code for localized RNN source estimation will be shared in a public repository before publication.

## 3. Results

### 3.1. Simulations

Estimated source signals from the RNN after training steps 1 and 2 for non-optimally and optimally oriented 12- and 54-source simulations are plotted in Supplementary Figures S6-S13. It can be seen that training step 2 produces more efficient, sparser source estimates, which were considered for all subsequent analyses unless stated otherwise.

Figure 3 presents examples of ground truth simulated source activations (vector magnitude) and results from four different source estimation methods for the optimally-oriented, 54-source simulation at select source indices. Comparable figures for the non-optimally and optimally oriented 12-source simulation, and the non-optimally oriented 54-source simulation, are provided in Supplementary Figures S14-S16, respectively. The BF method estimates noise present in the inactive source (index 0) and underestimates highly correlated bilateral sinusoidal activations at left (index 11) and right (index 15) temporal sources; this sinusoidal activation is also present in the frontal source (index 53) estimate. The BF method reproduces the Hann-windowed feature in the frontal source in simulated deviant trials from 0.2 to 0.4 s relatively well, although this appears to be slightly overestimated and misshapen. The MNE and dSPM methods also estimate signals on the inactive source, and accurately estimate bilateral sinusoidal and frontal Hann-windowed source activations; however, MNE slightly underestimates and dSPM slightly overestimates source amplitudes. The RNN method estimated less activity on the inactive source compared with the other methods, and predicted zero activity in the pre-stimulus baseline window. It also slightly underestimated the amplitudes of bilateral temporal and frontal source activations.

**Figure 3.**
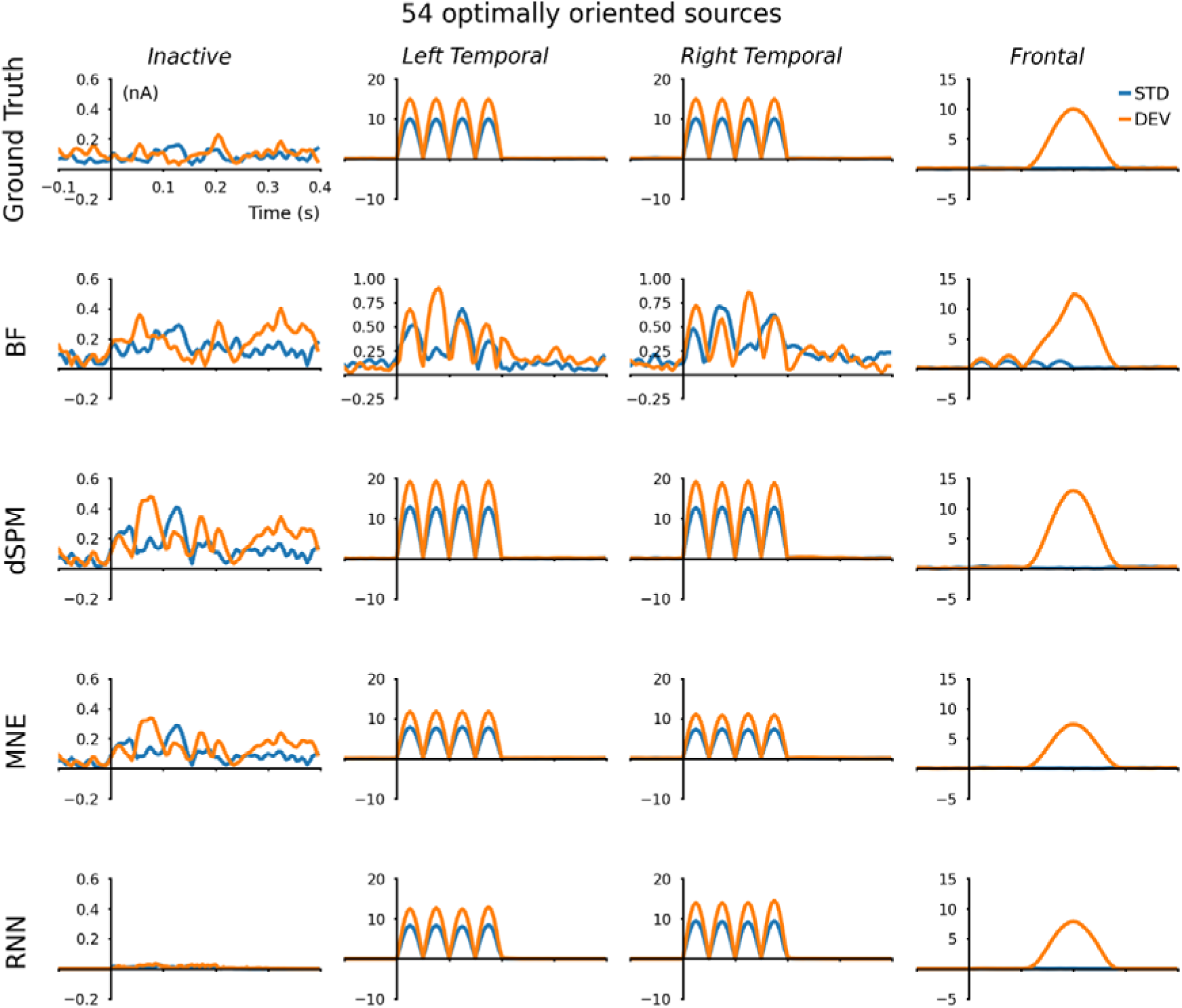
Source estimations from 54-source optimally oriented simulation. Ground truth simulated sources are plotted on the top row. Estimations from beamformer (BF), dSPM, MNE and RNN are plotted on the next four rows. The columns show an inactive source (index 1), left and right temporal sources (indices 11 and 15) and a frontal source (index 53). Blue traces are simulated standard activations, orange traces are simulated deviant activations. The inactive source is better estimated by the BF, dSPM and MNE estimators, with the RNN estimating tiny amplitude changes for this source after 0 s. Bilateral temporal sources are underestimated by the BF, overestimated by dSPM, and are comparatively well estimated by MNE and RNN. The frontal source is reproduced fairly-well by all estimators, although is slightly overestimated by BF and dSPM; the BF estimate of the frontal source activation is also contaminated with a sinusoid originating from temporal sources. Note that different y-axis scales have been used for some plots to improve visualization. Comparable results from 12 source and non-optimally oriented 54 source simulations are given in supplementary figures S14-S16.

Reconstructed ERFs from the four simulations are plotted in Figure 4. Optimally oriented sources produced higher amplitude simulated MEG. Also, 54-source simulations yielded higher amplitude simulated MEG than 12-source simulations, due to differences in the forward model. Unsurprisingly, because its computations do not consider the fitting of reconstructed ERFs to real ERFs, source estimations from the BF approach did not reproduce ERFs as well as the other methods, and it is evident that highly correlated activity has been underestimated [2]. Minimum norm methods (MNE and dSPM) produced reconstructed ERFs that were highly correlated with simulated ERFs. In time windows where only noise is present (i.e., <0 s for both, >0.2 s for standards, and >0.3 s for deviants), the MNE-reconstructed ERFs almost overlap the simulated ERFs. Although, in time windows of source activity, MNE-reconstructed ERFs have lower amplitude than the simulated ERFs. Source estimates from the RNN method were the closest to matching simulated ERFs.

**Figure 4.**
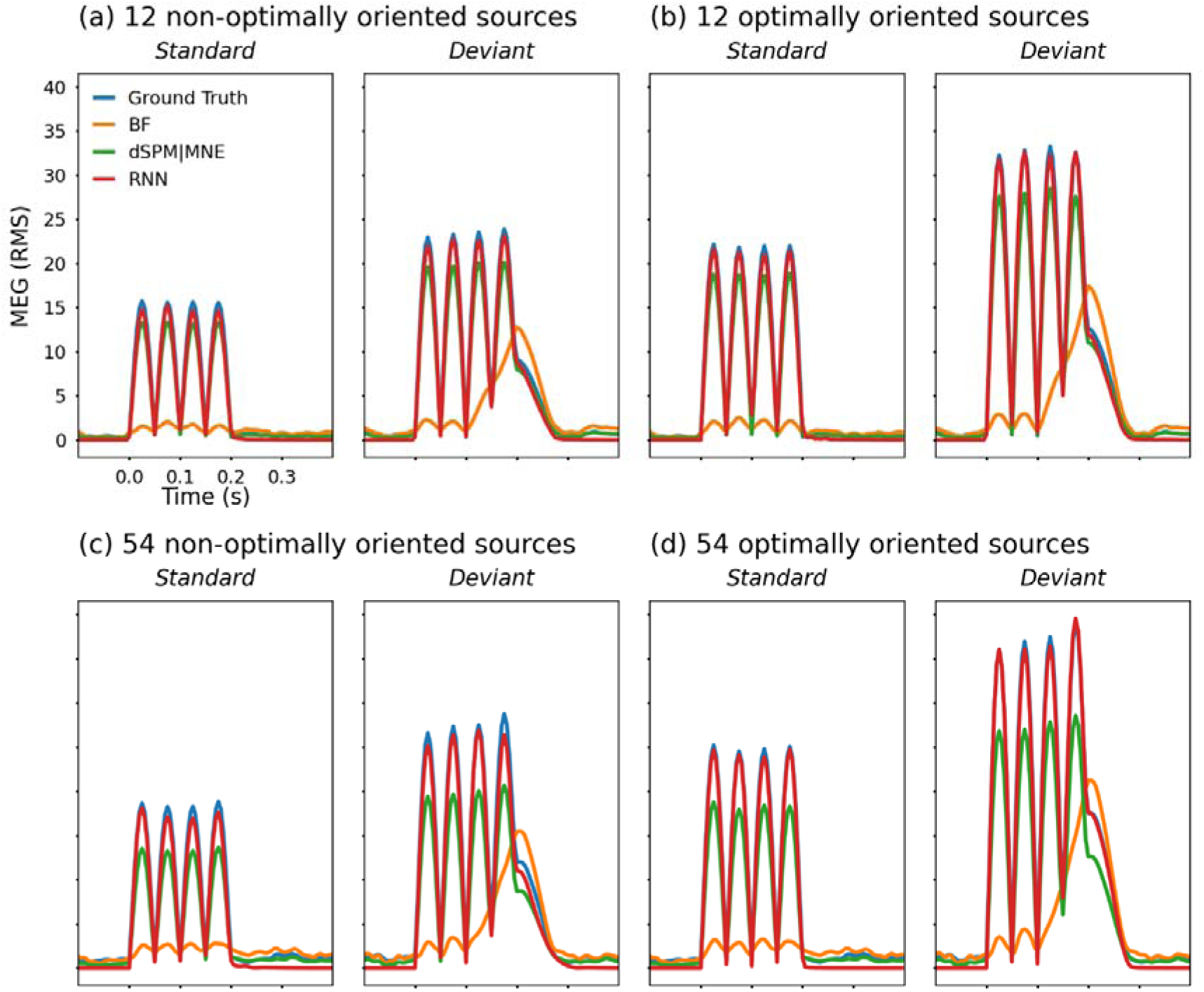
Simulated event-related fields and those reconstructed from estimated sources. Root-mean-squared amplitudes across 160 channels normalized to fT range are shown for (a) non-optimally, and (b) optimally oriented 12 source simulated standard and deviant signals; and also for (c) non-optimally, and (d) optimally-oriented 54 source simulated data. Optimal orientation of simulated source activations maximized the magnitude of resulting simulated MEG signals. Reconstructed ERFs from dSPM and MNE source estimates are overlapping and slightly under-estimate the simulated activity; BF, dSPM and MNE estimates track the noise pre- and post-stimulus, whereas the RNN does not reproduce the noise.

Analysis of source estimates and reconstructed ERFs from all of the simulations is provided in Table 1. The BF method consistently produced the lowest correlations, highest errors, and lowest SNRs between simulated and estimated sources and ERFs. The MNE and dSPM source estimates generally had the highest correlations with simulated source activity, although they were marginally surpassed by the RNN method for the 54-source optimally oriented simulation. Part of this high correlation and low error stem from MNE and dSPM estimates tracking the noise, as observed in Figures 3 and 4, which is undesirable. Source estimates using the RNN method effectively had infinite SNR because activity in the pre-stimulus baseline period was estimated to be zero, and otherwise achieved metrics comparable with MNE and dSPM estimates.

### 3.2. Real MEG data

Correlations among source estimates from five RNNs that were initialised with different random seeds are illustrated in Figure 5. Estimation layer outputs after training steps 1 and 2 are plotted separately. For the 12-source models, after step 1 there was an average correlation among source estimates of 0.681, which increased to 0.891 after training step 2. For the 54-source models, after step 1 the average correlation among source estimates was 0.486, which increased to 0.743 after step 2. Increasing average correlation among estimates after training step 2, which incorporated L1 regularization, indicates that the models converge towards a common solution. Source estimates from 54-source models correlating less than those from 12-source models reflects the larger space of possible solutions presented by a model with more parameters; despite this, the 54-source models reach fairly close alignment after training step 2. Source estimates from the RNN method taken forward for further analyses were obtained by averaging those from five seeded initialisations.

**Figure 5.**
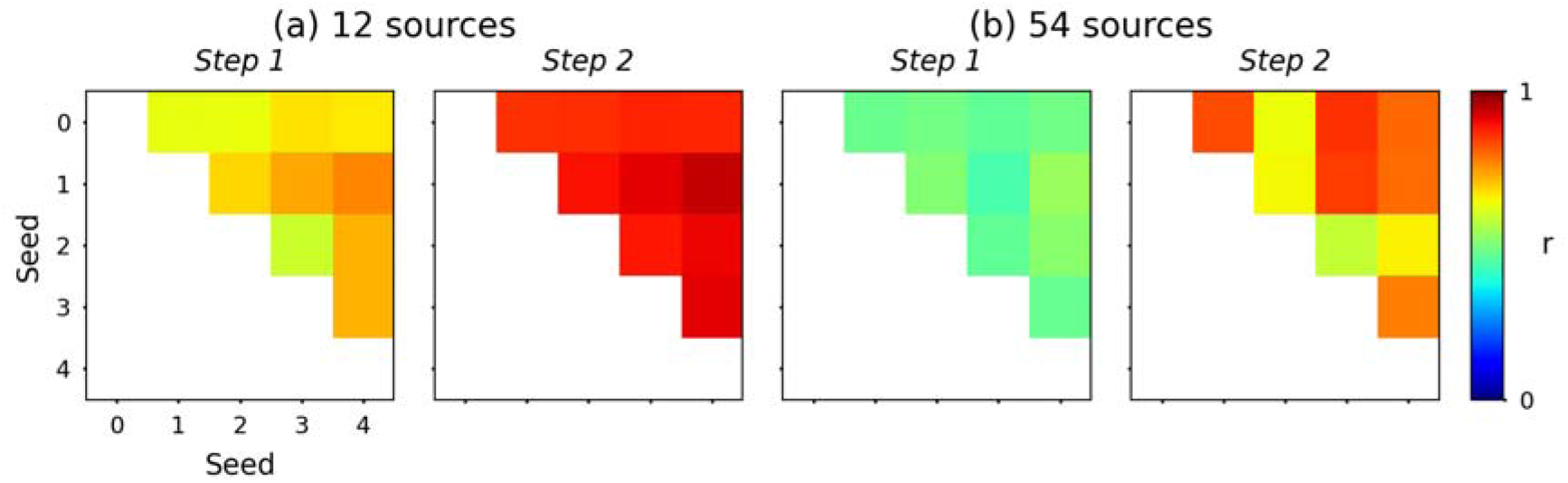
Evaluation of RNNs trained with different random seeds. Mean correlation among 12-source estimates from step 1 was 0.681 and step 2 was 0.891, and that among 54-source estimates from step 1 was 0.486 and step 2 was 0.743; correlations were calculated based on the resultant vector magnitude from x, y, and z components. Higher correlations between estimated sources following the second phase of training indicate that the models converge towards similar solutions.

Analysis of source estimation methods applied to real MEG data is provided in Table 2. Here we have only evaluated source estimates in terms of their signal-to-noise ratio; BF produced the lowest, then MNE, followed by dSPM-estimated sources, and the RNN method produced the highest SNR source estimates. Estimated sources were also evaluated for their ability to reconstruct the pattern of observed MEG signals. Here, again the BF approach underperformed relative to the other methods, producing lower correlations and higher errors between reconstructed and real ERFs. For the 12-source space, the RNN approach produced slightly higher correlation and slightly lower error between reconstructed and real ERFs than MNE and dSPM methods. For the 54-source space, this situation was reversed, and MNE and dSPM methods produced reconstructed ERFs with marginally higher correlation and lower error than the RNN approach. Estimated signals for each source location using the four estimation methods are provided in an online repository (https://osf.io/jxw2d/?view_only=cc654c4c6f4a464bbd08d860deb9c0a4).

The top ten sources based on various ranking methods are given in Table 3. There is some consistency among these rankings for different source estimation techniques. The MNE and dSPM methods produce identical results with sources ranked by SNR; perhaps unsurprisingly because dSPM estimates are scaled copies of MNE esitmates. For the 12-source estimates, indices 10, 8, 9, 3 and 4 tend to be highly ranked, thus may be considered as the most relevant sources underlying the observed pattern of MEG signals. For the 54-source estimates, indices 32, 46, 15, 27 and 28 tend to be ranked highly, although there is more variability in these rankings.

Source signals estimated using the RNN method are plotted in Figure 6 for the 12-source and Figure 7 for the 54-source forward models. By incrementally turning on these units in order from highest to lowest SNR, their relative contributions to reconstructed ERFs were calculated in terms of correlation with real ERFs. As shown in Figure 8, this analysis suggests that all of the sources in the 12-source model were required to achieve performance achieved using 20/54 (37%) of the sources in the 54-source model. As such, the 54-source model provided redundancy to derive a sparse pattern of source signals; in contrast, the 12-source model could not achieve a sparse representation, and instead generated signals across all sources. This lack of redundancy in the 12-source space could explain estimated signals of questionable plausibility at source indices 6, 8 and 11 in Figure 6. Inspecting Figure 7 for comparison, there are fewer large amplitude source estimates along the midline, as these tend to be concentrated towards temporal sites. The overall pattern of estimated sources signals from the 54-source model is consistent principally with temporal generators (indices 10, 15, 23, 27, 28, 32, 46 and 49) and contributions to the deviant response from right frontal-temporal (indices 35 and 52) and orbitofrontal (index 19) source locations.

**Figure 6.**
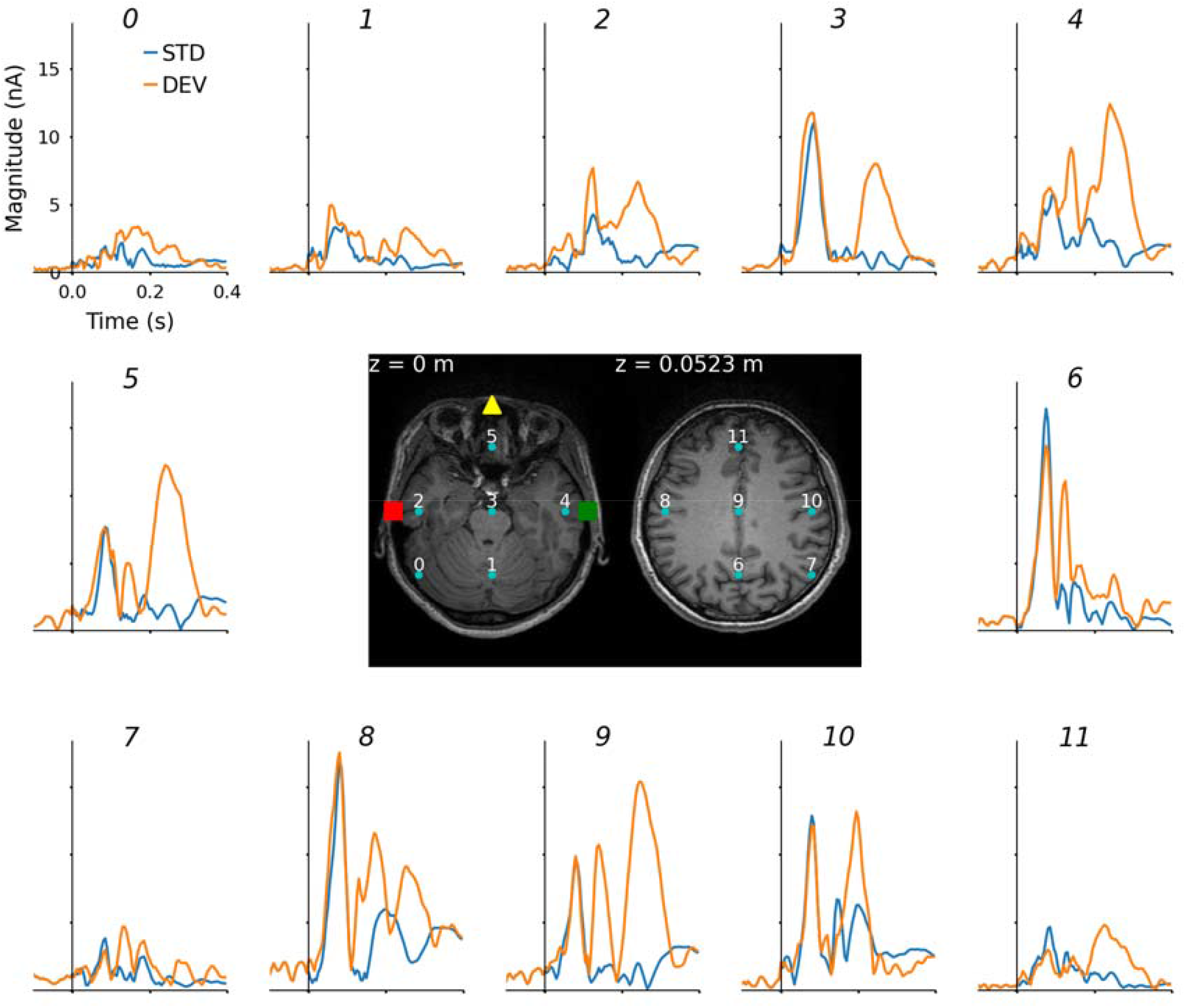
RNN-estimated source activations from real MEG data with 12-source source space. Vector magnitudes from three directions are plotted. Indices noted above each graph correspond to source locations annotated on MRI slices in the centre. For a comparison between source activations estimated using different methods, please see the figures provided in the “12 sources” folder of the online repository. The nasion is identified with a yellow triangle, and left and right preauricular areas are identified with red and green squares.

**Figure 7.**
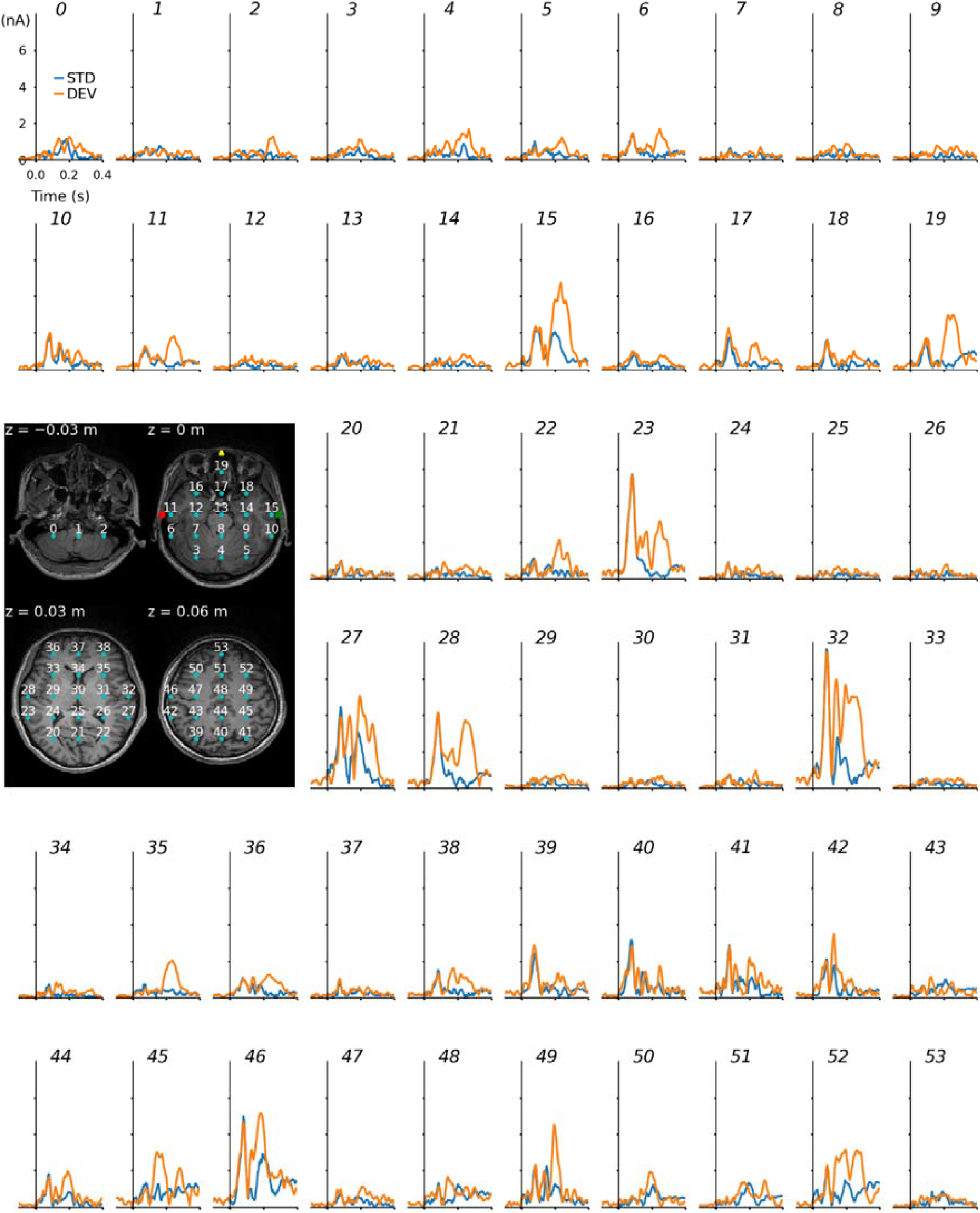
RNN-estimated source activations from real MEG data with 54-source source space. Vector magnitudes from three directions are plotted. Indices noted above each panel correspond with source locations identified in the MRI slices shown at the centre-left of the figure. Please see the figures provided within the “54 sources” folder in the online repository for a comparison between source activations estimated using different methods. The nasion is identified with a yellow triangle, and left and right preauricular areas are identified with red and green squares.

**Figure 8.**
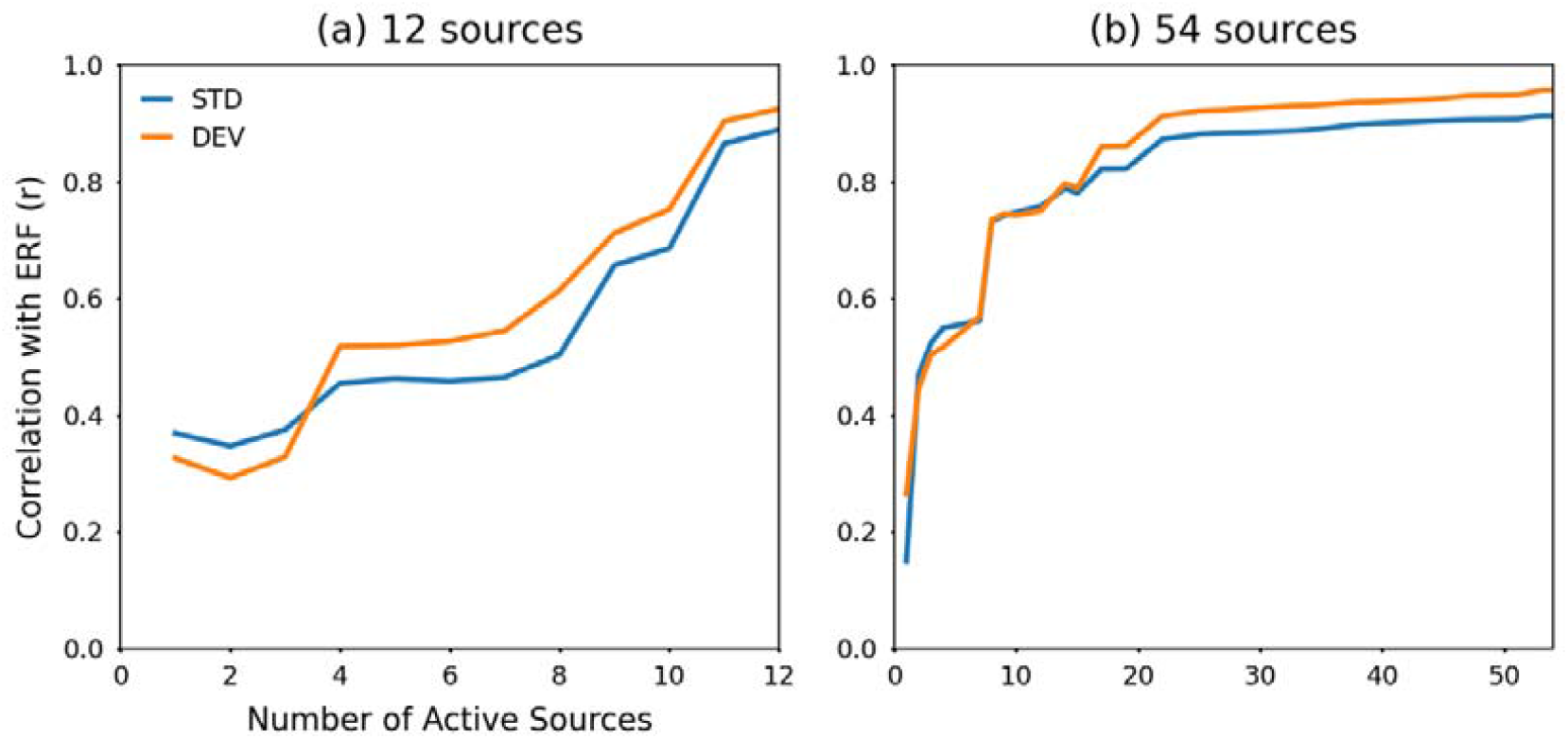
Correlation between real and reconstructed ERFs as RNN sources are turned on. Sources were incrementally activated based on their SNR, from highest to lowest. The 12-source source space (a) with 11 sources active reaches comparable levels of correlation as the 54-source source space (b) with 20 sources active. This observation suggests that the 54-source source space has more redundancy with which to fit the ERFs, leading to a sparser pattern of estimated source signals, as evidenced by comparing Figures 5 and 6.

In Figure 9, real ERFs are plotted alongside those reconstructed from source estimates produced using different methods. The BF approach leads to underestimated ERF amplitudes in the time window of the initial auditory response (i.e., from 0 to 0.1 s post-stimulus); although the later portion in which differences between standard and deviant responses emerge is reproduced comparatively well when the 54-source forward model was used. The MNE and dSPM reconstructed ERFs, and those from RNN-estimated sources, similarly reproduced ERFs for both 12- and 54-source forward models.

**Figure 9.**
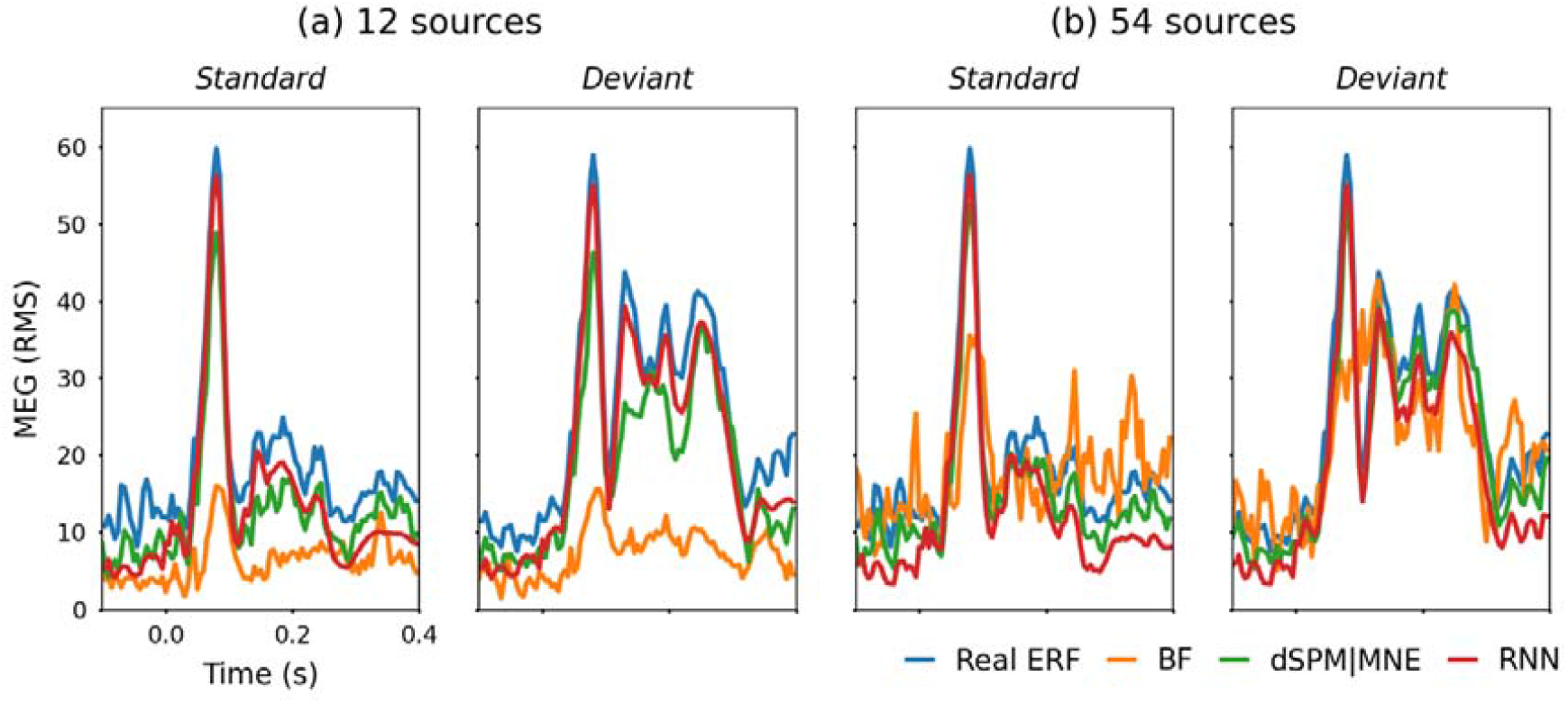
Comparison between real ERFs and those reconstructed from source activations estimated using different methods. Results from (a) 12 source and (b) 54 source spaces are plotted. The RNN approach yielded the highest SNR in both situations and the highest correlation with real ERFs when 12 sources were used. In the 54-source source space, dSPM, MNE and RNN approaches correlated similarly with real ERFs.

## 4. Discussion

From the simulation results in Table 1, it is apparent that optimal source orientation led to better performance for the RNN approach regarding similarity between estimated and simulated source signals. In contrast, the other methods were influenced less by the orientation of simulated sources. The BF approach was inferior to the others across all metrics used, presumably due to its acknowledged limitations [2,26]. Although several beamforming modifications have been proposed to compensate for these limitations which may result in better performance [5,26]. The RNN method produced infinite output SNR because it did not track the noise; in contrast, other methods estimated source activity pre-stimulus and also for inactive sources that only contained noise. Otherwise, estimated source signals from MNE, dSPM and RNN methods were reasonably comparable, as seen in Figure 3, as were the ERFs reconstructed from these estimates (Figure 4).

Five seeded initialisations of the RNN model were applied to real MEG data. These produced source estimates that became increasingly correlated after the second training step, where L1 regularization was applied to the source estimation layer (Figure 5). The number of model initialisations could have been increased to obtain a more robust average estimate. However, this would proportionally increase the computational demands that were already considerably greater than the other methods. To illustrate the magnitude of the difference in computational requirements, consider that BF, MNE and dSPM source estimates were computed within seconds. In contrast, two training phases for an RNN were completed within several hours. Thus, with regard to computation time, the other methods are evidently superior to the RNN method.

It is more challenging to evaluate estimated source signals for genuine MEG data than it is to evaluate estimates of simulated data. In terms of output SNR, which we can consider, the RNN method outperformed the others. Reconstructed ERFs from MNE, dSPM and RNN methods were comparable, although the RNN had higher SNR, as noted in Table 2 and seen from ERFs in Figure 9. However, comparing estimated source signals with real source signals is unfeasible because the latter are unknown. Alternatively, we can compare rankings and correlation among source signals estimated by each method. Please view the source signals estimated by different methods shared in an online repository (https://osf.io/jxw2d/?view_only=cc654c4c6f4a464bbd08d860deb9c0a4). Consistency among source estimates and their rankings, for example indices 10 and 32 for 12- and 54-source spaces, respectively (Table 3), provide strong evidence that these sources contribute significantly to the observed pattern of MEG signals. Both sources are located in the right temporal lobe, consistent with expectations for auditory ERFs. The overall pattern of estimated sources from the RNN plotted in Figures 6 and 7 is consistent with bilateral temporal, right-frontal and orbitofrontal cortex generators, which generally concurs with the literature on MMNm [14].

The 12-source space arguably does not contain enough sources for the RNN to derive a usefully sparse representation. Consider that the RNN required all of the sources in the 12-source space to be active to fit the output labels with r ≈ 0.9, whereas equivalent performance with the 54-source space was reached with 20/54 active sources (Figure 8). Redundancy within the source space allows L1 regularization to encourage the RNN in training step 2 to find an efficient solution to fit the labels, which drives down the amplitude of sources that contribute less effectively to the observed MEG. The resulting signals might be similar to those produced by minimum-norm methods that incorporate the L1 norm as an additional constraint [28–30]. Visual inspection of the source estimates in Figures 6 and 7 also leads us to conclude that the 54-source space produces more reasonable results.

Whilst the 54-source space seems preferable to the 12-source space, a case could be made that it is still too sparse, and a denser source space would improve localization of source estimates. There are a couple of points to consider concerning increasing the source density when using the RNN approach. Firstly, increasing the number of units in the model architecture to design an appropriately sized source estimation layer would further increase its computational requirements, which may or may not become prohibitive, depending on the number of sources and available computing resources. Secondly, relatively sparse source spaces are less susceptible to forward model errors introduced by small changes in head position [20]. Provided that head movements are less than the volumetric source grid spacing, we can justifiably assume that the forward model (and lead field matrix) remains constant. However, a static forward model becomes a less viable assumption for dense source spaces where head movements may be greater than the grid spacing. Furthermore, in situations where forward model errors can be large, e.g., studies in young children who might have difficulty keeping their head still [31], it may be necessary to track head position and update the forward model continuously. Although it is unclear precisely how the RNN method of source estimation might be adapted to compensate for continual changes in the forward model, we can speculate on devising a training scheme where lead field weights are synchronised and paired with data labels (single-trial MEG) to allow the network to represent these changes.

Several other methods using deep neural networks have been proposed for source localization and signal estimation [20,32–35]. Pantazis and Adler developed deep and convolutional neural networks (DNNs and CNNs) for predicting underlying source coordinates from one time-sample or a brief snippet of MEG signals [20]. This approach is reportedly tolerant to forward model errors and potentially suitable for real-time localization of a predefined number of sources; although in practice, real-time coordinates might not provide meaningful information without also being able to observe estimated signals from those locations. Hecker et al. also used a CNN to localize sources from a single time-sample of EEG data and reported that their model outperformed other state-of-the-art methods [34]; however, it is unclear from their paper whether this approach is capable of estimating source signals that underlie event-related potentials. Dinh et al. used a type of RNN consisting of Long Short-Term Memory (LSTM) cells to correct estimates generated by minimum-norm algorithms [33]. The resulting corrected source estimates were more precise than those estimated from minimum-norm methods, highlighting the sequence-modelling benefits of RNNs. Sun et al. trained a DNN on simulated data from neural mass models combined with a forward volume conduction model to estimate source activity underlying EEG [32]. They validated this approach using visual and somatosensory evoked responses and intracranial measurements of epileptiform neural activity. Liang et al. proposed combining sparse Bayesian learning with a DNN and evaluated their approach using simulations [35]. Collectively, these studies highlight the promise of applying deep learning methods for estimating electromagnetic source signals.

Recurrent neural networks offer a considerable degree of flexibility that may set them apart from the aforementioned deep-learning approaches. They have previously been used to simulate changes in auditory evoked responses that occur between states [9], which could be adapted to estimate how state changes influence the dynamics of underlying source signals. Simulations can also be performed to examine how the output of an RNN changes as its input representations are varied; for example, manipulating represented stimulus duration, frequency or intensity [10], altering representations of sound loudness [11], or changing input images to models of visually-evoked potentials [13]. Similar manipulations could be applied to models developed for localized source estimation to investigate what aspects of the modelling paradigm influence estimated source signals. Conversely, incorporating L1 regularization into RNNs trained for non-localized source estimation could enforce sparser representations that might be easier to evaluate and interpret. It is also possible to modify the loss function used for optimizing the RNN to encourage desirable behaviour, for instance by adding a noise-to-signal ratio component to maximize output SNR. Although only applied to data from a single subject in this study, RNNs could be used for group-level inference by applying to each subject individually, or applying to a grand-averaged dataset then using transfer learning to fit data from individual subjects and quantifying changes between sets of estimated sources.

## 5. Conclusion

We have presented a way to estimate electromagnetic source signals underlying event-related neural activity using RNNs. In this method, the RNN is initially trained to minimize mean-squared-error loss between its outputs and MEG labels, then fine-tuned with L1 regularization applied to its final hidden layer, in both cases with the output layer weights fixed as the lead field matrix from the forward model. After the second training phase, estimated source signals are obtained from the final hidden layer activations. This method has been validated using simulations and compares favourably with other established methods. It clearly outperformed beamformer, MNE and dSPM methods in terms of output SNR, and performed comparably in terms of correlation and error between estimated and simulated source activity. When applied to real MEG data recorded during an auditory oddball experiment, the RNN method estimated source signals that generally concurred with expected brain locations based on the literature, yielding patterns of neural activity consistent with temporal, right frontal-temporal, and orbitofrontal generators. The main drawback of the RNN approach is its computational demands, which are several orders of magnitude greater than the other methods. Nevertheless, its advantages (extendable network architecture, flexible training paradigm, capacity for running simulations, transfer learning, reverse-engineering, etc.) make this an invaluable tool for studying computational transformations between sensory/cognitive events and neural activity [36–38].

## Acknowledgements

JAO was sponsored to train with PFS by the International Brain Research Organisation (IBRO) Asia Pacific Regional Committee (APRC) with a 2022 Exchange Fellowship. This work was also supported by King Mongkut’s Institute of Technology Ladkrabang (grant number KREF186607), an Australian Research Council Discovery Project (grant number DP170103148), and the Australian National Imaging Facility.

## Data availability

Data and code from this study will be shared publicly via an Open Science Foundation repository (https://osf.io/jxw2d/?view_only=cc654c4c6f4a464bbd08d860deb9c0a4).

## Ethical review

The experimental protocol was reviewed and approved by the Human Research Ethics Committee at Macquarie University.

## Author Contributions (CRediT Statement)

JAO: Conceptualization, Methodology, Software, Formal Analysis, Data Curation, Writing - original draft, Writing - reviewing and editing, Visualization, Funding acquisition.

JDZ: Conceptualization, Methodology, Software, Data Curation, Writing - reviewing and editing.

PFS: Conceptualization, Resources, Writing - reviewing and editing, Supervision, Funding acquisition.

