## Supplemental Figures for "Localized estimation of electromagnetic sources underlying event-related fields using recurrent neural networks"

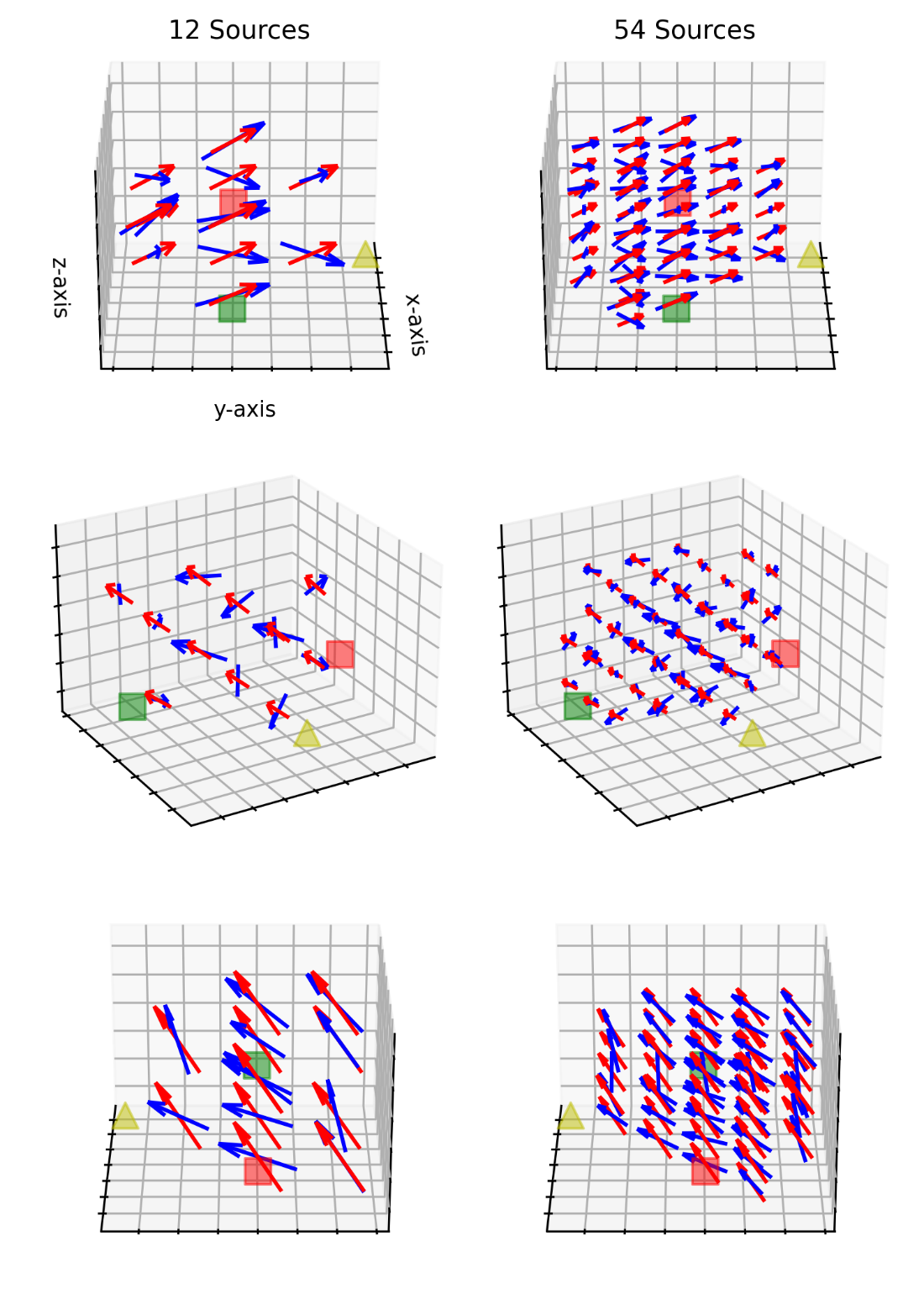

**Figure S1** Illustration of volumetric source spaces with non-optimal and optimal dipole orientations. Source spaces with 12 and 54 sources are shown on left and right columns, with different view angles (0°, 60° and 180°) shown across three rows. Red and blue arrows represent non-optimal and optimal orientations of resultant simulated source activations. The nasion is identified with a yellow triangle; left and right preauricular areas are denoted with red and green squares. Non-optimally oriented vectors are all pointing in the same direction, whereas optimally oriented vectors are directed parallel to the plane of the closest sensors, reflecting a magnetic field orthogonal to the nearest pickup coil. The length of all arrows is equal to the grid spacing, and they are centred at their respective source location.

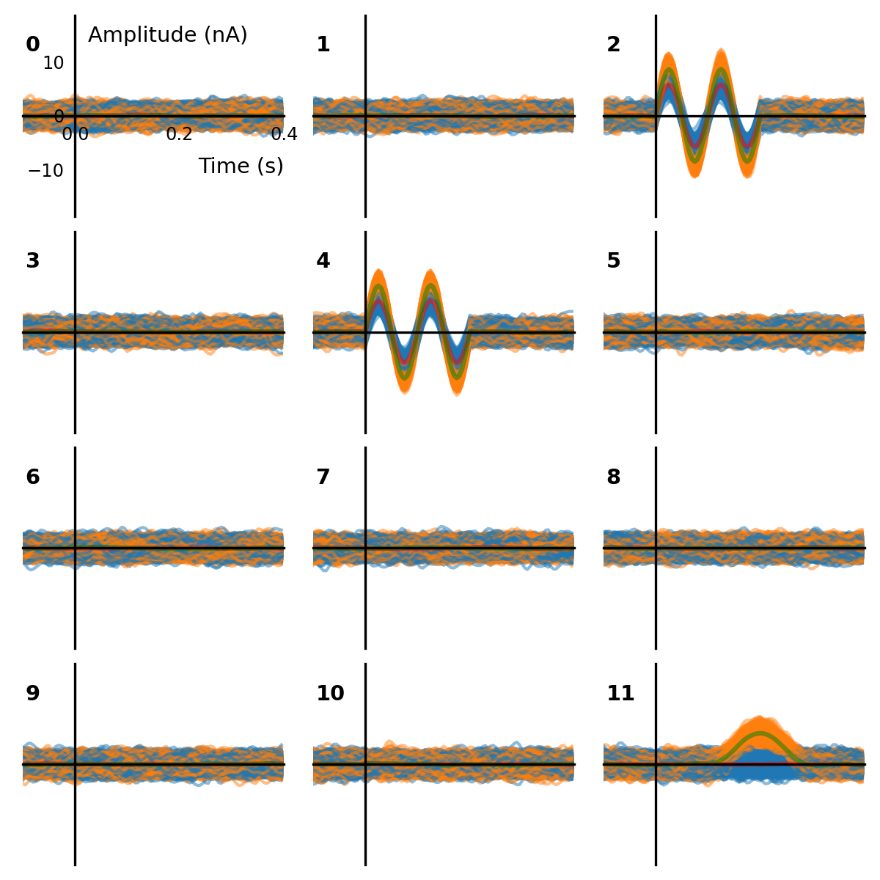

**Figure S2**  Non-optimally oriented 12-source simulations. Simulated “standard” trials are plotted in blue, and stimulated “deviant” trials are plotted in orange. Source indices 2 and 4 were chosen for simulating bilateral sinusoids that differ in amplitude between standard (10 nA peak) and deviant (15 nA peak) trials. Source index 11 was selected for a frontal time-delayed Hann-window activation only in deviant trials with peak amplitude of 10 nA. Average source activations for the simulated standard trials are plotted in red and for the simulated deviant trials are plotted in green. Vector magnitude in x-, y- and z- directions are identical in this non=optimally oriented case.

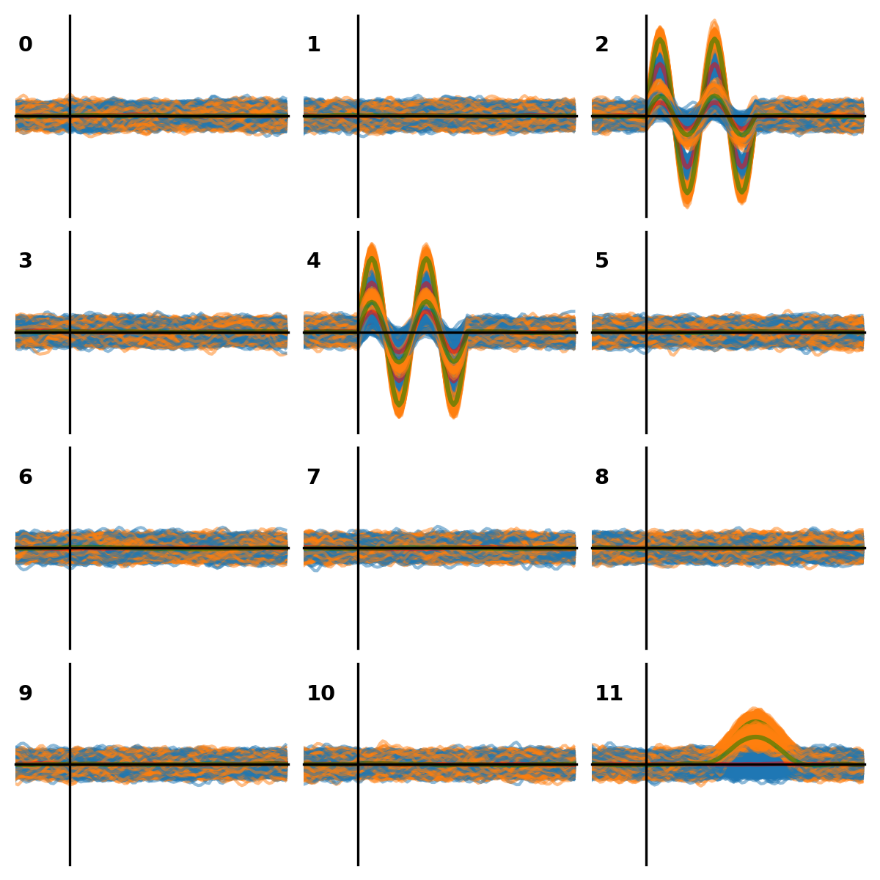

**Figure S3**  Optimally oriented 12-source simulations. The x- and y-axes, and line colour scheme are the same as those in Figure S1. In this optimally oriented case, resultant vector magnitudes remained the same (indices 2 and 4 peak amplitudes of 10 nA for simulated standards and 15 nA for simulated deviants; index 11 peak amplitude of 10 nA for simulated deviants), however, magnitudes in x-, y-, and z-directions are unequal, reflecting an optimum weighting derived from the lead field matrix.

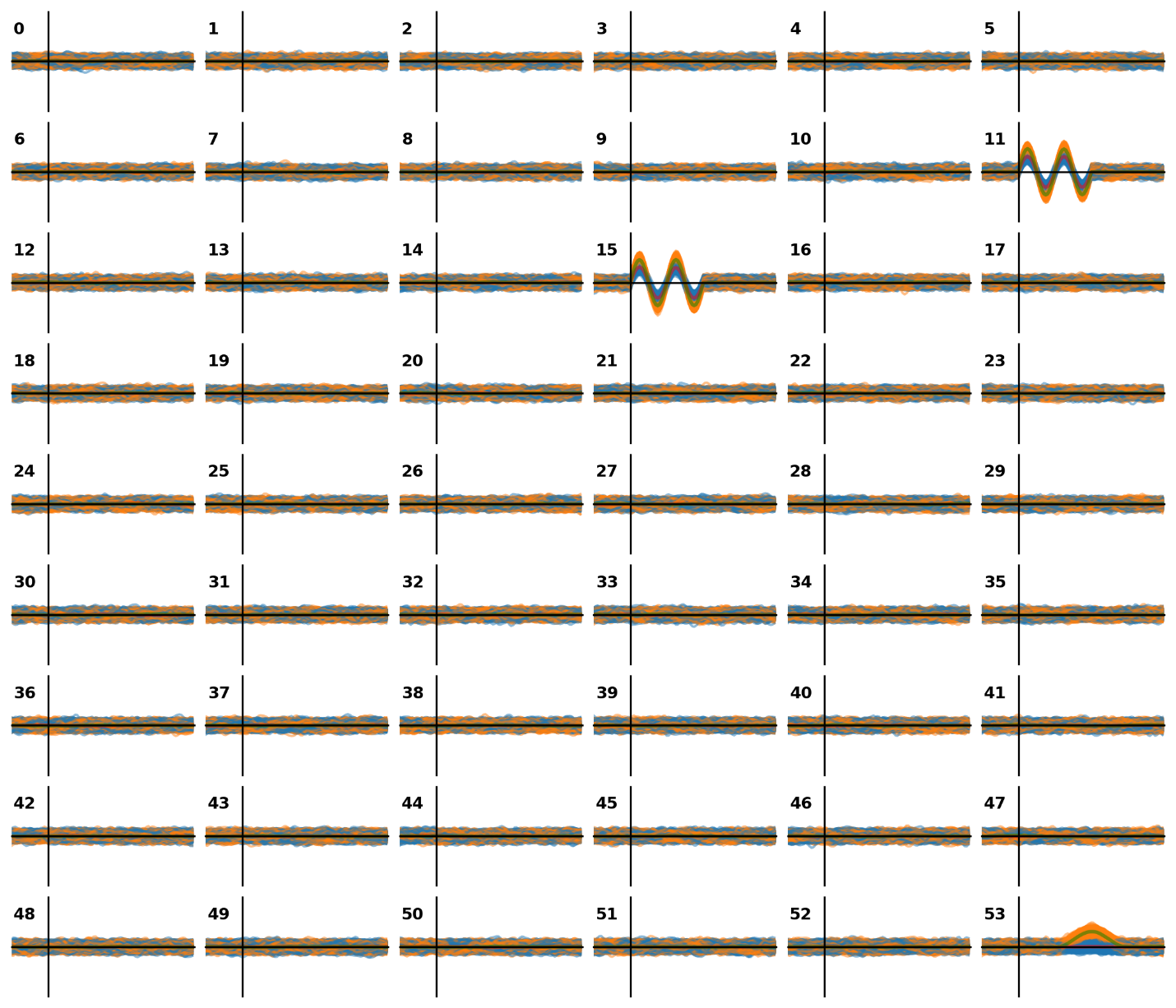

**Figure S4**  Non-optimally oriented 54-source simulations. The x- and y-axes, and line colour scheme are the same as those in Figures S1 and S2. Source indices 11 and 15 were bilaterally located in the temporal lobes, source index 53 was located in the frontal lobe.

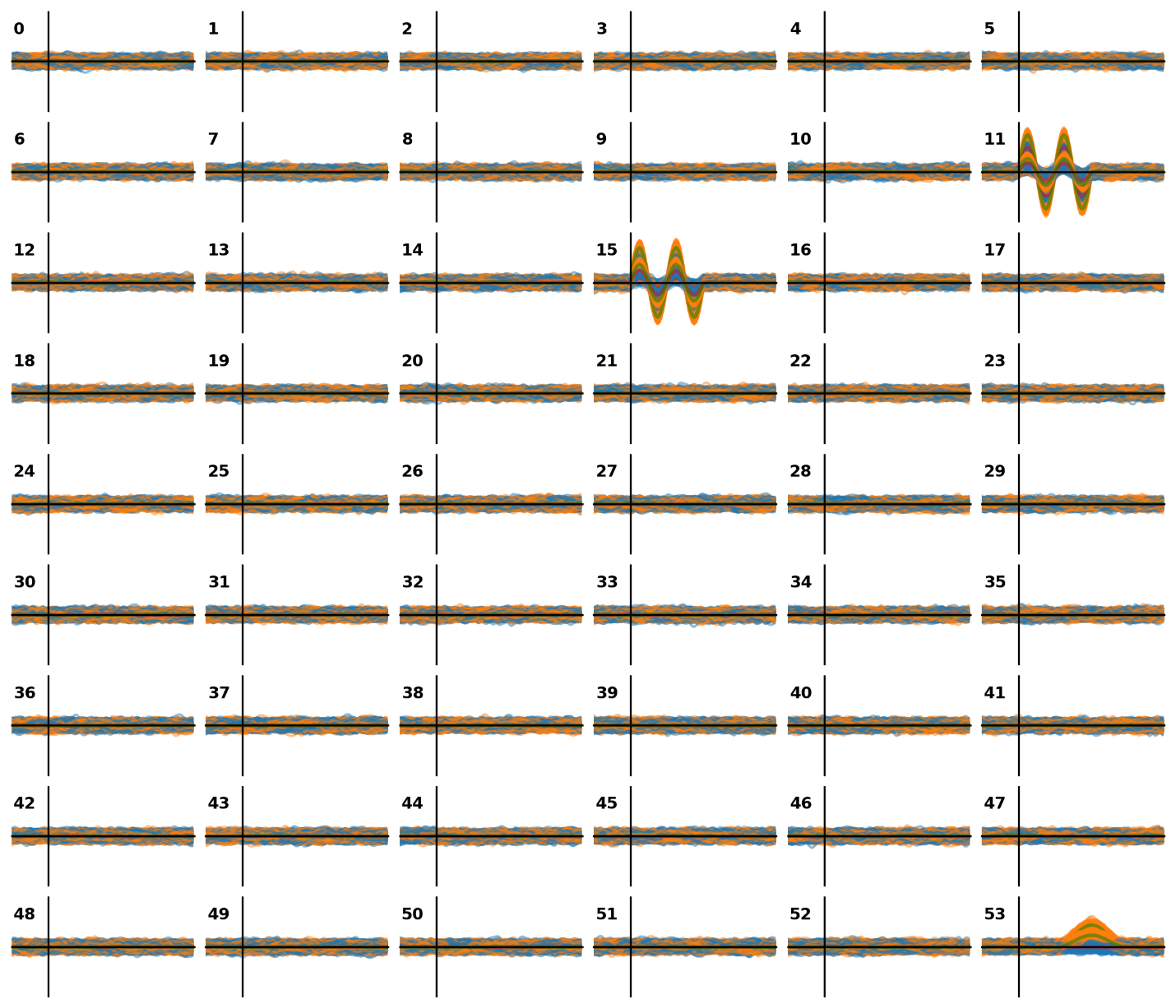

**Figure S5** Optimally oriented 54-source simulations. The x- and y-axes, and line colour scheme are the same as those in Figures S1-S3. Optimal orientation is achieved from varying the x-, y-, and z-components of simulated sources.

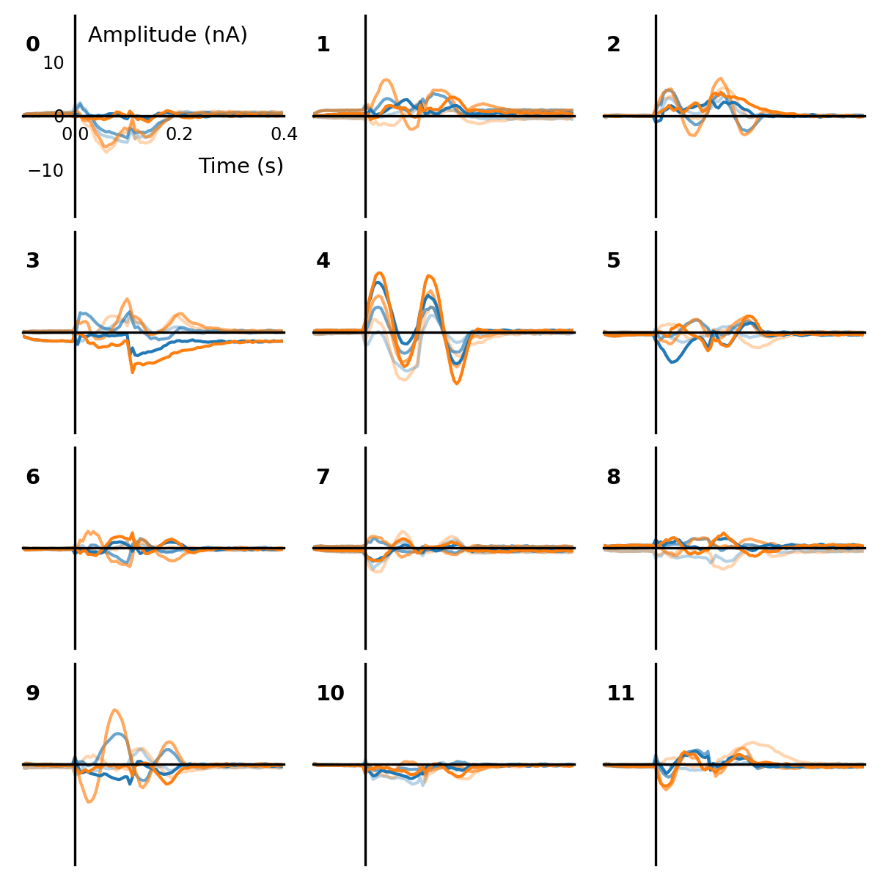

**Figure S6** Non-optimally oriented 12-source source space, RNN step 1. Standard in blue, deviant in orange.

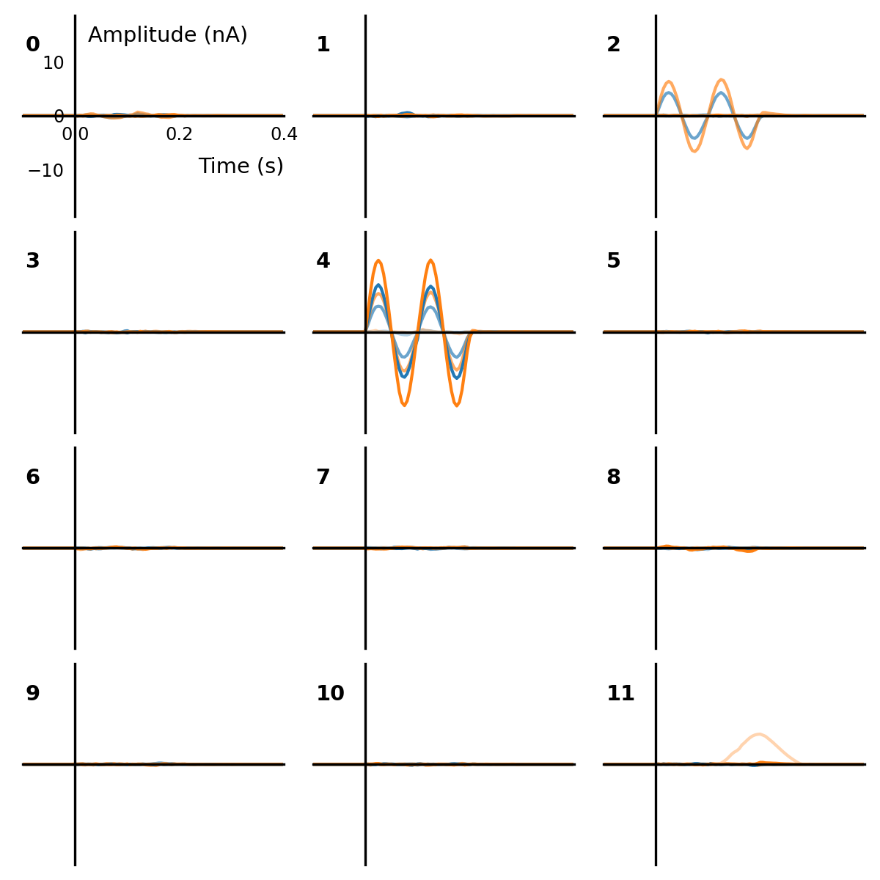

**Figure S7** Non-optimally oriented 12-source source space, RNN step 2. Standard in blue, deviant in orange.

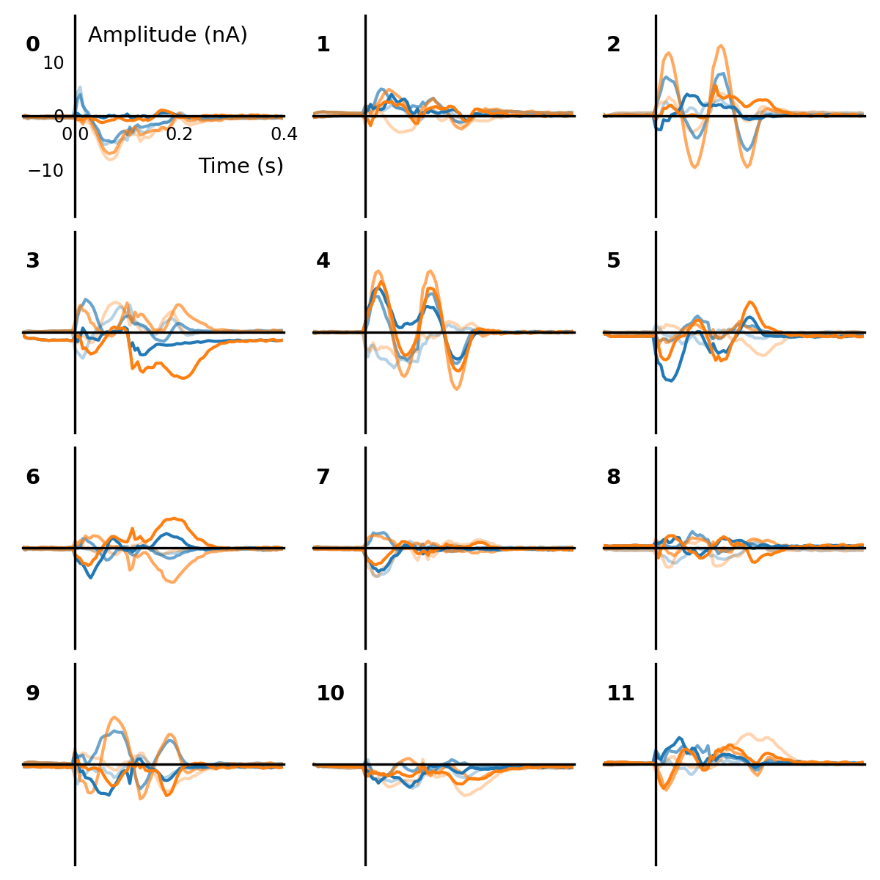

**Figure S8** Optimally oriented 12-source source space, RNN step 1. Standard in blue, deviant in orange.

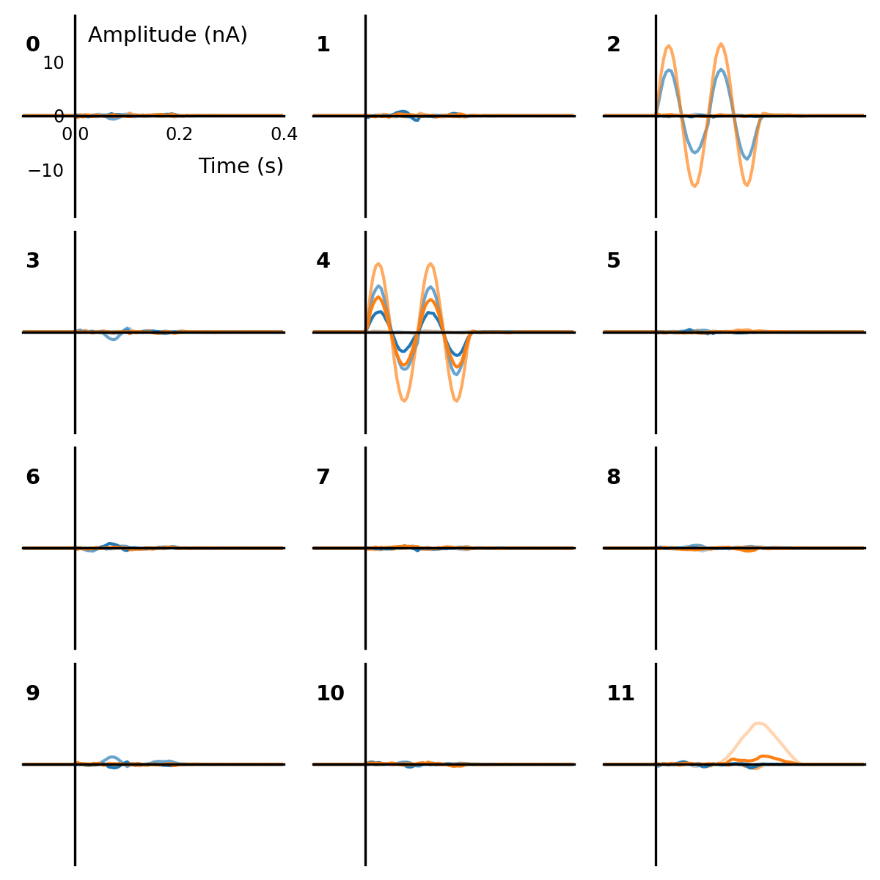

**Figure S9** Optimally oriented 12-source source space, RNN step 2. Standard in blue, deviant in orange.

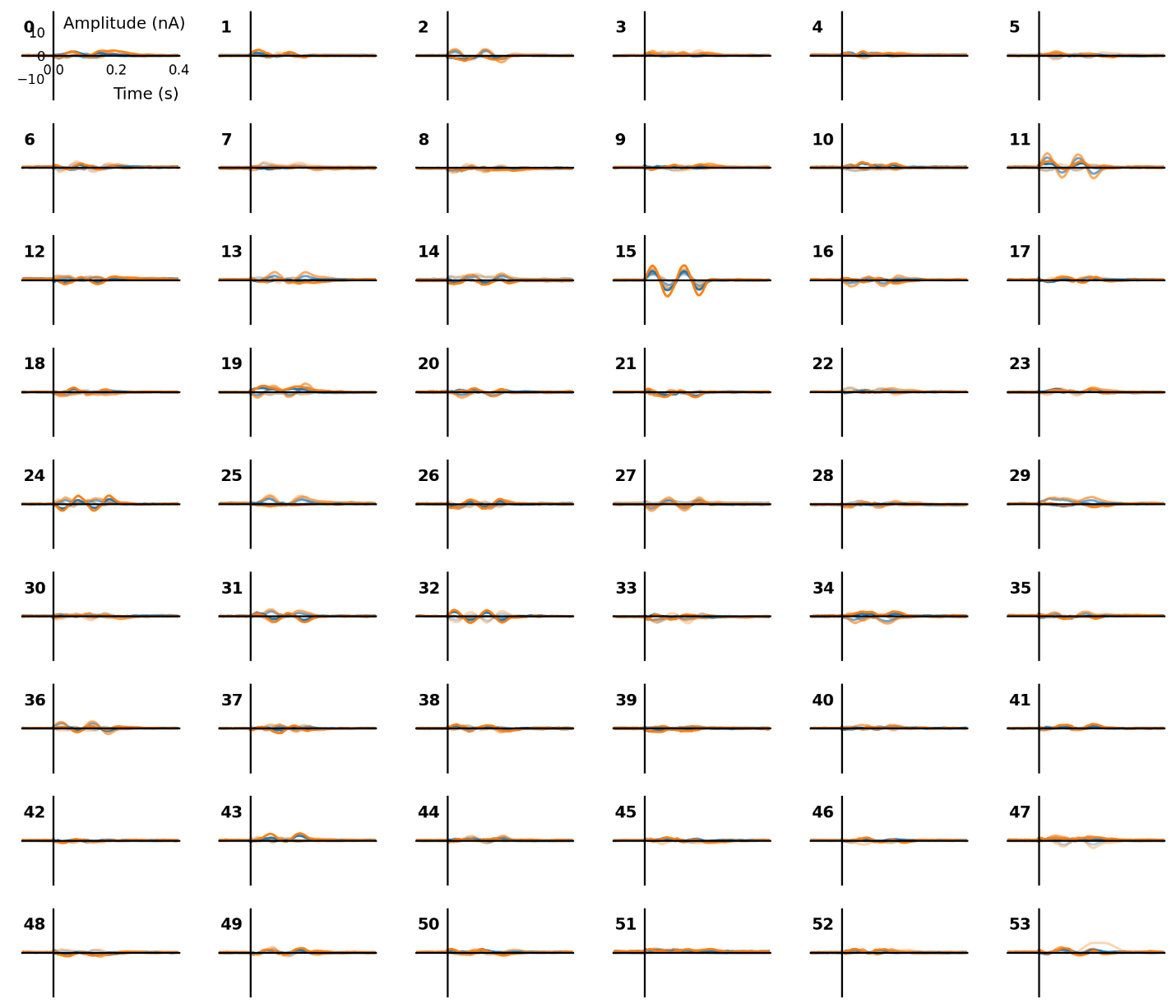

**Figure S10** Non-optimally oriented 54-source source space simulation, RNN step 1 result. Standard in blue, deviant in orange.

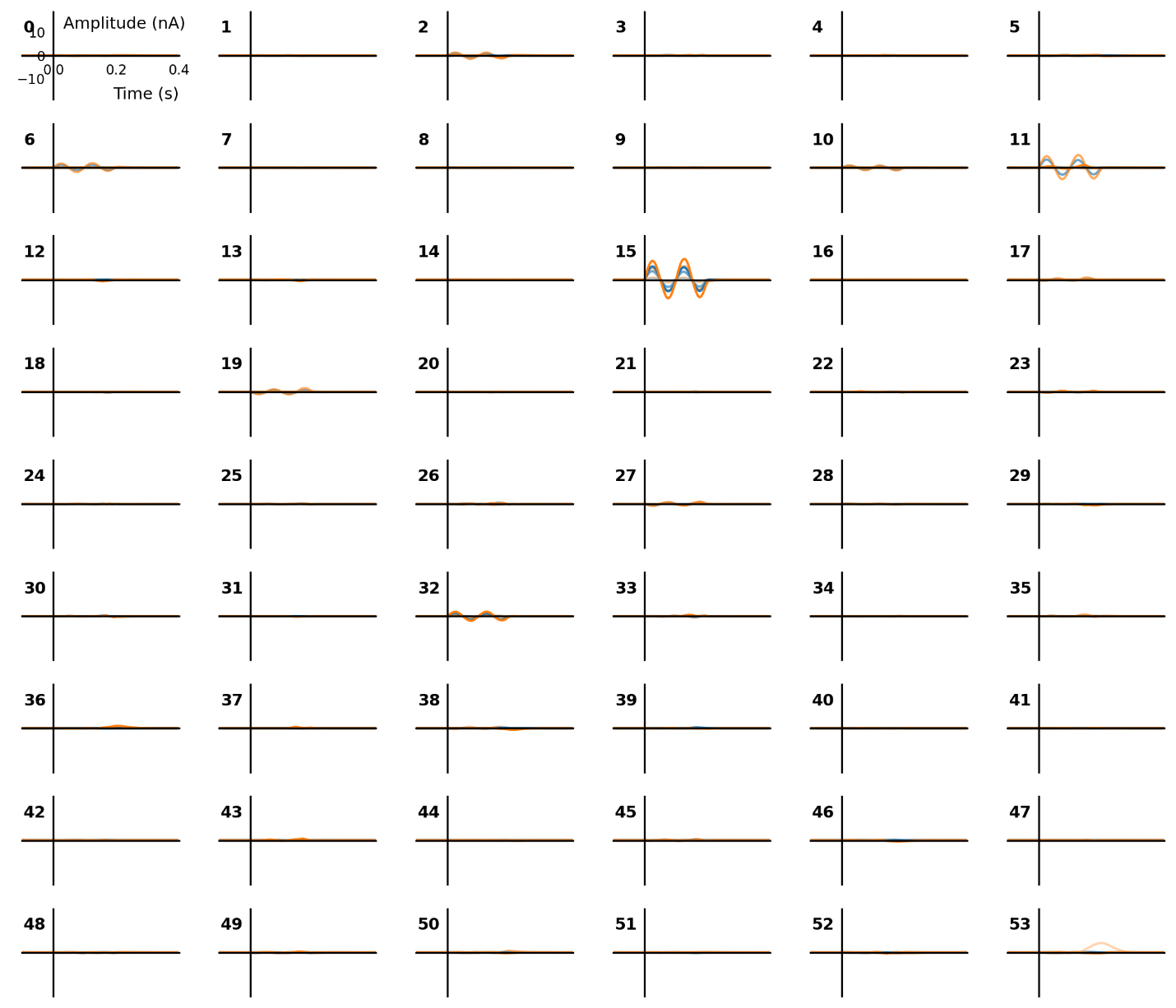

**Figure S11** Non-optimally oriented 54-source source space simulation, RNN step 2 result. Standard in blue, deviant in orange.

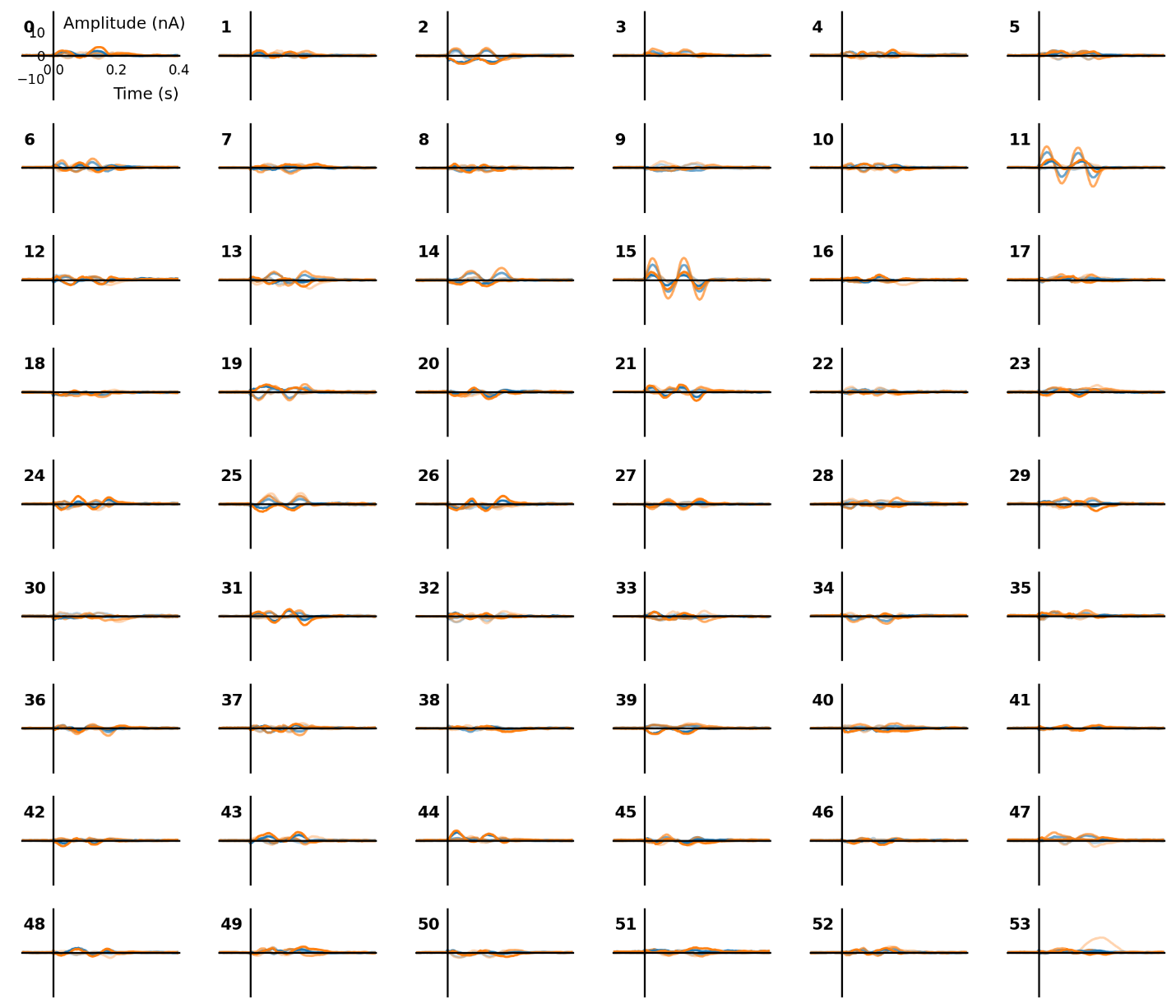

**Figure S12** Optimally oriented 54-source source space simulation, RNN step 1 result. Standard in blue, deviant in orange.

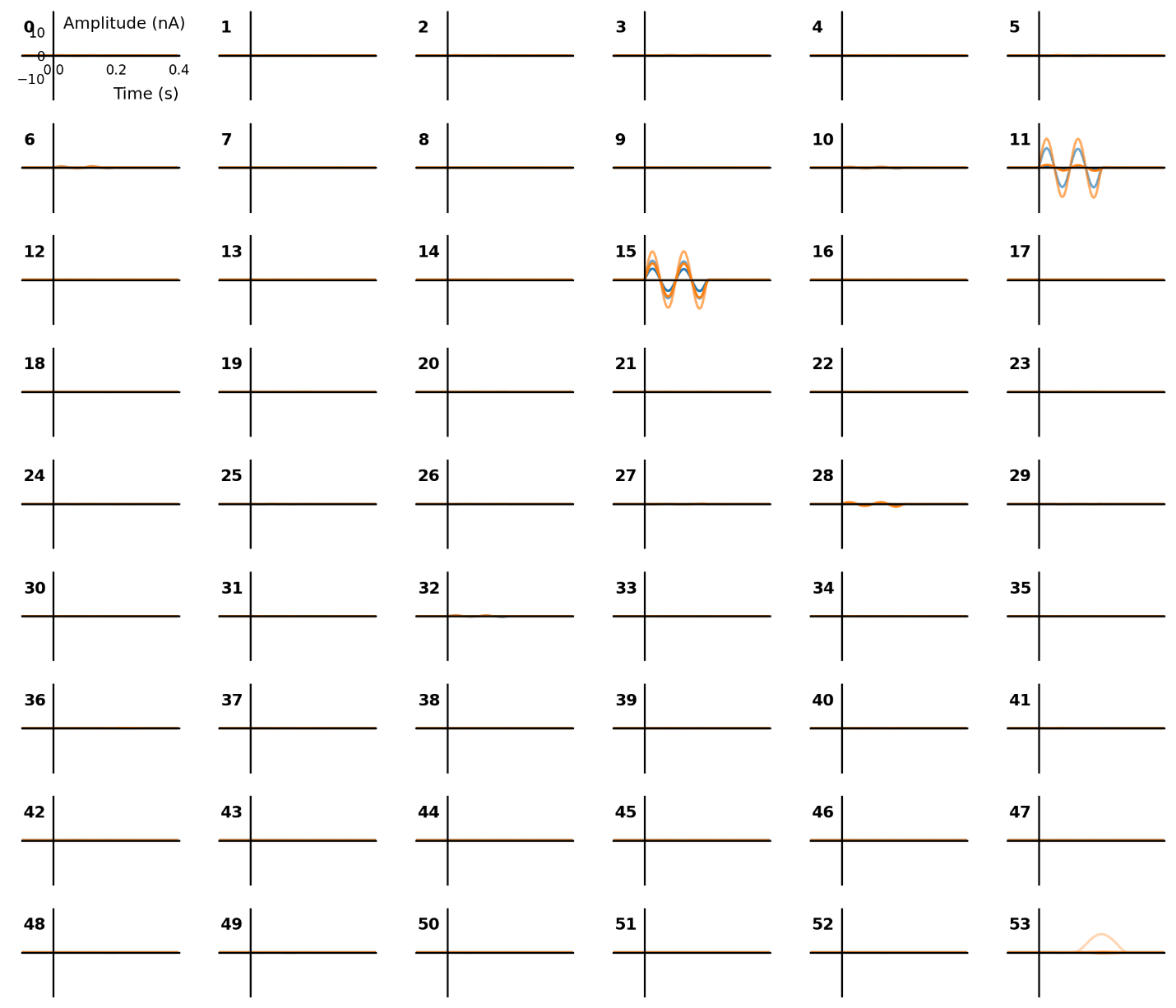

**Figure S13** Optimally oriented 54-source source space simulation, RNN step 2 result. Standard in blue, deviant in orange.

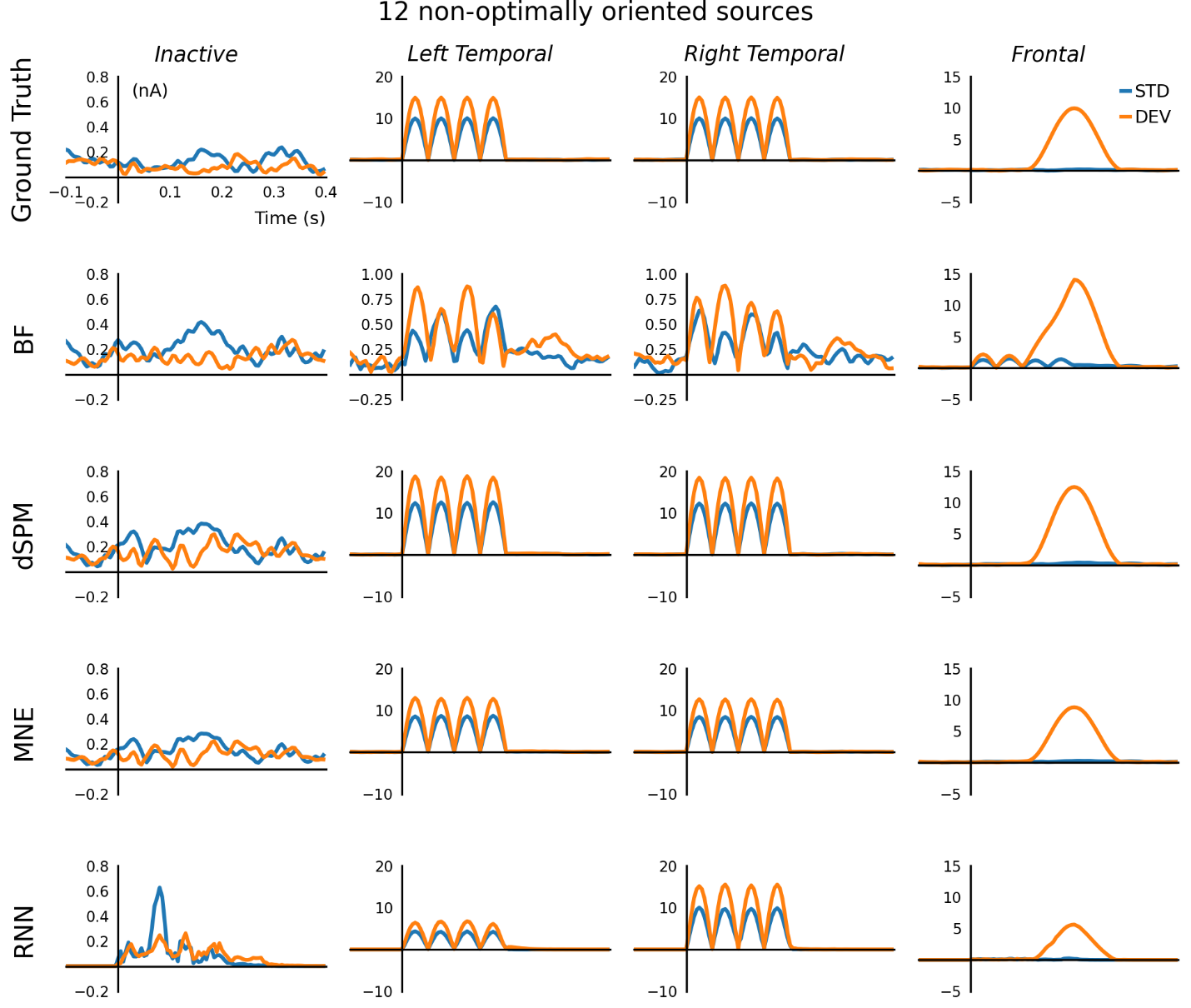

**Figure S14** Comparison of estimated sources from simulation with 12 non-optimally oriented sources. Vector magnitudes from three spatial directions are plotted.

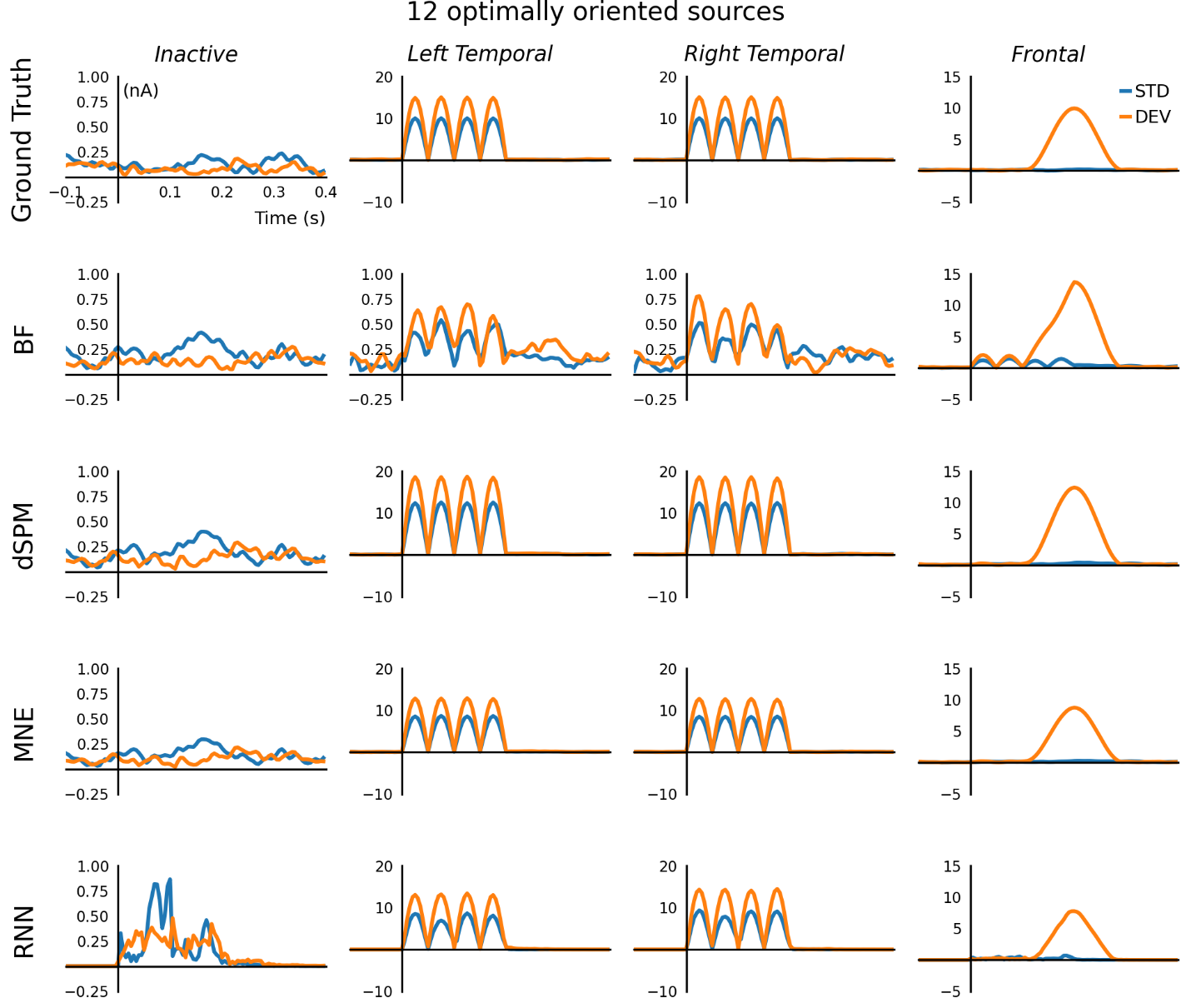

**Figure S15** Comparison of estimated sources from simulation with 12 optimally oriented sources. Vector magnitudes from three spatial directions are plotted.

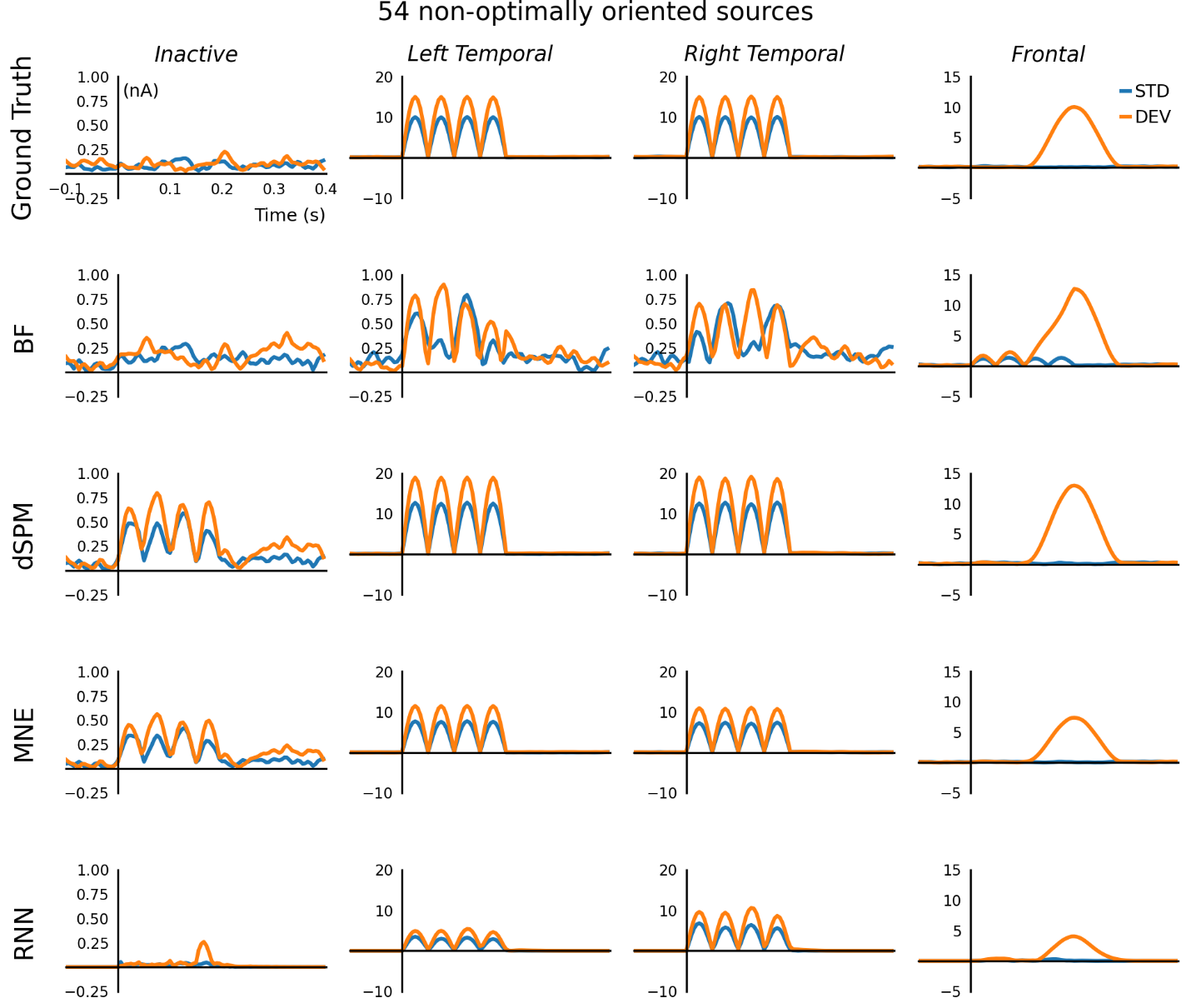

**Figure S16** Comparison of estimated sources from simulation with 54 non-optimally oriented sources. Vector magnitudes from three spatial directions are plotted.
